# Somatic *DDX41* mutations confer neomorphic splicing activity in the pathogenesis of myelodysplastic neoplasms

**DOI:** 10.64898/2026.09.23.753593

**Authors:** Rasika Venkataraman, Noemi Puccio, Tun-Yun Hsueh, Elizabeth A. Bonner, Fu-Yen Chang, Erica A. Arriaga-Gomez, Axia Song, Takero Miyagawa, Jahan Rahman, Oluwanifemi Akinwande, Sayantani Sinha, Ettaib El Marabti, Robert S. Welner, Rui Lu, P. Brent Ferrell, Martin Carroll, Jacob S. Appelbaum, Derek L. Stirewalt, Omar Abdel-Wahab, Anthony Rongvaux, Xueyan Chen, Kikkeri N. Naresh, Sebastian M. Fica, Stanley C. Lee

## Abstract

Germline mutations in the RNA helicase *DDX41* are the most common genetic predisposition to myelodysplastic syndromes (MDS) and acute myeloid leukemia (AML), representing 5-7% of AML. Over 70% of patients acquire a somatic mutation at specific hotspots (*R525H/G530D*) in the helicase domain of DDX41 in *trans*, which is linked to disease progression. DDX41 has diverse cellular functions, including RNA splicing, ribosome biogenesis, R-loop resolution and inflammation. However, the mechanisms by which somatic *DDX41* mutations drive MDS/AML pathogenesis are yet to be determined. Here, we systematically evaluated the function of pathogenic, missense germline and somatic *DDX41* mutations. We report that *DDX41* somatic mutations are gain of function alleles and fail to rescue AML cell growth, distinct from germline *DDX41* mutations. *DDX41* somatic mutations R525H and G530D drive widespread splicing changes in isogenic AML cell lines, consistent in genetically engineered murine bone marrow HSPCs and CD34^+^ cells from *DDX41*-mutant MDS/AML patients. Mechanistically, DDX41-R525H exhibits increased RNA binding affinity compared to wildtype, with aberrantly spliced RNA targets overlapping with enriched binding. Strikingly, aberrant splicing of SEPTIN7 is enriched exclusively in CD34^+^ HSPCs of patients harboring both germline and somatic *DDX41* mutations but absent in samples with germline mutation alone or lacking *DDX41* mutations altogether. SEPTIN7 mis-splicing introduces a premature stop codon resulting in its downregulation at the protein level. SEPTIN7 is a known regulator of cytokinesis, HSC polarity and repopulation capacity. Our results identify a novel mis-spliced candidate, SEPTIN7, whose deregulation implicates a mechanism by which the *DDX41* somatic mutations promote dysplastic hematopoiesis.

## INTRODUCTION

Germline mutations in the gene *DDX41* are the most common genetic predisposition to myeloid neoplasms including myelodysplastic syndromes (MDS), myelodysplastic/myeloproliferative neoplasms (MDS/MPN) and acute myeloid leukemia (AML), representing 5-7% of adult patients with myeloid neoplasms^1–3^. Patients with germline heterozygous mutations in *DDX41* generally present with clinical symptoms of cytopenia and myeloid malignancy at a median age of 70 years^1,4^, similar to sporadic disease. These patients present with low leukocyte count, hypocellular bone marrow, and are predominantly male^5,6^. In addition to having relatively normal cytogenetic profiles, *DDX41* mutant patients have distinct co-mutation patterns relative to *DDX41* wildtype (WT) patients. Over 60% of patients with germline *DDX41* mutations acquire a somatic missense mutation in *trans* enriched at hotspots in the helicase domain (p.R525H or p.G530D) of DDX41, which is associated with progression to AML^2,3,7^. While *DDX41* germline mutant patients often acquire additional MDS/AML-associated driver mutations, such as *ASXL1*, *SRSF2*, *TP53*, and *TET2*, only *CUX1* and *GNAS* mutations are significantly enriched in *DDX41* biallelic mutant patients relative to *DDX41* WT patients^2,8,9^.

DDX41 is a multifunctional DEAD-box RNA helicase and has been shown to participate in diverse cellular processes, including nucleic acid sensing and innate immune signaling^10–13^, ribosome biogenesis and translation control^7,14,15^, R-loop homeostasis and genome stability^13,16–18^ and pre-mRNA splicing^3,17,19–21^. In addition, *Ddx41* is essential for embryonic development and hematopoiesis. Conditional knockout of *Ddx41* in mice caused hematopoietic stem cell (HSC) defects prenatally, most severely of the myeloid lineage, and *Ddx41* knockout mice failed to survive postnataly^22^. Homozygous deletion of *Ddx41* or bialleic-R525H-null mutations impaired hematopoiesis, resulting in bone marrow failure^14,23^. Despite considerable progress, the specific downstream effectors linking mutant DDX41 to hematopoietic dysfunction and leukemogenesis remain undefined.

Structurally, DDX41 contains a set of conserved domains characteristic of the DEAD-box helicase family: an N-terminal region that is predicted to be intrinsically disordered, a DEADc domain that binds nucleic acid and ATP, a HELICc domain that mediates ATP hydrolysis, and a C-terminal region containing a short zinc finger domain^3,24^. Germline *DDX41* mutations are heterozygous; two-thirds of which are truncating (e.g., p.D140fs, p.A500fs) or start-loss (p.M1I) variants and are predicted to result in loss of function^1,3^. Non-truncating germline variants are enriched in the DEAD domain of DDX41, and among these, p.E256K, p.Y259C, and p.S363del are likely pathogenic^2^. In contrast, somatic *DDX41* missense mutations are found primarily at two hotspots within the HELICc domain (p.R525H or p.G530D), which diminish ATPase and helicase activity^2,3,7^. How germline loss of one *DDX41* allele combined with a somatic HELICc mutation in the other allele drives MDS and AML pathogenesis remains poorly understood.

The ATPase and helicase activities of DEAH-box and DEAD-box protein families drive the conformational rearrangements within the spliceosome that are required for pre-mRNA splicing^25^. Additionally, DDX41, along with other DEAD-box RNA helicases, has been identified as a component of the catalytically active spliceosome^19,26^. Given that recurrent mutations in other spliceosomal genes^27,28^ (e.g. *SF3B1, SRSF2, U2AF1* and *ZRSR2*) are established drivers of MDS/AML, we hypothesized that somatic *DDX41* mutations, such as p.R525H, which reduce ATPase activity^7^, would impair RNA processing and cause aberrant splicing of key regulators of HSC biology, ultimately leading to dysplastic hematopoiesis.

In this study, we systematically examined RNA splicing in both *DDX41* germline-and somatic-mutant AML cells using a genetic complementation system. We defined and validated a splicing signature specific to *DDX41* somatic variants with germline lesions using a combination of isogenic human AML cell lines, disease-relevant mouse models, and bone marrow specimens from patients with MDS and AML. These findings identify potential downstream effectors linking somatic *DDX41* mutations to the pathogenesis of myeloid malignancy.

## METHODS

### Antibodies and oligos

All primers, short hairpin RNA (shRNA) and antibody information used in this study are available in Supplementary Tables S1 and S2, respectively.

### Cell culture

MOLM-13 cells were maintained in base media of RPMI 1640 (Gibco #22400089) and 293T cells in DMEM (Gibco #11965092), supplemented with penicillin-streptomycin (100 U/mL, Gibco #15140122), GlutaMAX (1%, Gibco #35050061), sodium pyruvate (1 mM, Gibco #11360070) and 10% heat-inactivated fetal bovine serum and maintained in a 37°C, 5% CO_2_ incubator.

### DDX41 constructs

cDNA expressing DDX41 wildtype or mutant coding sequence were cloned into pRSC32 vector backbone, received from Dr. Hans-Peter Kiem (Fred Hutchinson Cancer Center). Short-hairpin RNA targeting DDX41 (sh.DDX41) was designed using the SplashRNA algorithm^29^ and cloned into the pLT3GEPIR^30^ vector (a gift from Dr. Johannes Zuber, Addgene Plasmid #111177).

### Lentivirus production

Lentiviral particles were produced using 293T cells seeded in 10 cm tissue culture dishes at a density of 3.4×10^6^ cells in 10 mL of complete DMEM. 24 hours post seeding, cells were co-transfected with packaging plasmids pVSVG (4 μg) and psPAX2 (6 μg) along with expression plasmids (10 μg) using the CalPhos™ Mammalian Transfection Kit (Takara #631312). Viral mix was removed 16h post transfection, cells were gently rinsed with 1X PBS (Gibco) and replaced with complete RPMI (see above). Viral supernatant was collected 48 and 72 hours post transfection, filtered of cell debris using Millex™-GV Filter Unit (Millipore Sigma, SLGVM33RS), snap frozen and stored in −80°C.

### Transduction and cell sorting

MOLM-13 cells were transduced with a 1:1 dilution of viral supernatant in the presence of polybrene (5 μg/mL; Millipore #TR-1003-G) for 48h. For shRNA transduction, cells were selected with 2 μg/mL of puromycin dihydrochloride (Gibco #A1113803) post transduction and treated with 500 ng/mL of doxycycline for 48-72 hours to induce the shRNA. For cDNA transduction, GFP^+^ cells were sorted four days post transduction using the BD FACSymphony™ S6 Cell Sorter (BD Biosciences).

### Cell proliferation assay

MOLM-13 cells expressing doxycycline inducible sh.DDX41 and re-expressing DDX41 WT or mutant cDNA were sorted for GFP positivity. On recovery post sort, 60,000 cells were plated in 3.5 mL of complete RPMI with 2 μg/mL puromycin, with and without 500 ng/mL doxycycline in a 12 well culture plate. Live cell number was counted on days 3, 4, 5 and 7 post doxycycline treatment using the ViCell Blu cell counter (Beckman Coulter).

### Western blotting

Cells were lysed in IP-lysis buffer (Invitrogen, #87787) supplemented with protease inhibitors (ThermoFisher, #78446) and incubated on ice for 30 minutes. Lysates were centrifuged at 13,000rpm for 15 minutes to collect supernatants and protein concentrations were normalized using a Pierce BCA protein assay kit (Thermo Fisher Scientific #23227). 30 μg of protein lysate was mixed with Pierce™ Lane Marker Reducing Sample Buffer (Thermo Fisher, #39000), boiled for 5 minutes, loaded in pre-cast SDS acrylamide gels (BioRad #4561084) and run at 90V. Protein was transferred to PVDF membranes using the Trans-Blot® Turbo™ Transfer System (Bio-Rad #1704150EDU) following manufacturers recommendation. Membranes were blocked in Tris-buffered saline with 0.05% Tween-20 (TBS-T) and 5% milk for 1 hour at room temperature and immunoblotted with primary antibodies overnight at 4°C. Membranes were washed 3 times with TBS-T and incubated for 1 hour at room temperature with secondary antibodies conjugated to horseradish peroxidase. Blots were imaged using Clarity™ (BioRad, 1705060) or Clarity Max™ Western ECL Substrate, (BioRad, 1705062) using the ChemiDoc MP Imaging System (BioRad).

### RT-PCR

200 ng – 1 μg of Trizol extracted RNA was used to make cDNA using Verso cDNA Synthesis Kit (ThermoFisher, #AB1453B). PCR was performed with isoform specific primers (listed in Supplemental Table S1) using GoTAQ per manufacturer’s instructions (Promega, #M7123).

### RNA-sequencing

MOLM-13 cells expressing doxycycline inducible sh.DDX41 and re-expressing DDX41 WT or mutant cDNA were sorted for GFP positivity. On recovery post sort, 1 million cells were plated in 6 mL of complete RPMI with 2 μg/mL puromycin, with 1 μg/mL doxycycline in a 6 well plate. Samples were harvested 72 hours post doxycycline treatment for RNA extraction. RNA-sequencing libraries were prepared using NEBNext Poly(A) mRNA Magnetic Isolation Module (NEB #E7490) and NEBNext® Ultra™ II Directional RNA Library Prep Kit (NEB #E7760L). Bulk RNA-seq samples were sequenced using the NovaSeq X plus (Illumina) platform, generating 150-base pair paired-end reads.

### Bioinformatic analysis

FASTQ files were quality-checked using FastQC v0.12.0 and trimmed with Trim Galore v0.6.10 to remove adapter sequences and reads with Phred scores below 15. STAR v2.7.0a was used to align the FASTQ files to the hg38 reference genome with two-pass mode enabled. The resulting SAM files were sorted using the sort function from Samtools v1.19.2 and indexed using the index function to generate BAM index files. Gene expression was quantified using featureCounts v2.0.6 (Subread package), configured to count fragments assigned to exon features, and summarized by gene name based on the GENCODE v45 gene annotations. Differential gene expression analysis was performed using DESeq2 v1.42.0. Alternative splicing event detection was performed using rMATS v4.2.0 with paired-end mode, allowing for novel splice sites, variable read lengths, and soft-clipped reads. For each splicing comparison, statistically significant splicing events were defined as having: (i) an absolute inclusion level difference greater than 20%, (ii) a False Discovery Rate (FDR) less than 0.05, and (iii) a mean junction read coverage of at least 50 reads across replicates. For visualization of splicing events, BAM files were converted to bigWig format using bamCoverage from deepTools v3.5.4, with signal normalized using reads per kilobase per million mapped reads (RPKM). Read coverage tracks at statistically significant splicing events were then generated using pyGenomeTracks v3.8, enabling visual inspection of splicing patterns across conditions.

### Enhanced Crosslinking and Immunoprecipitation (eCLIP)

eCLIP was performed by Eclipse Bioinnovations Inc (San Diego) according to the published single-end seCLIP protocol^31^ with the following modifications. MOLM-13 cells were UV crosslinked at 400 mJoules/cm2 with 254 nm radiation, pelleted, and stored at −80°C until use. Cells were lysed using approximately 750 μL of eCLIP lysis mix with 3 μL Proteinase Inhibitor Cocktail and 10 μL of Murine RNase Inhibitor. Samples were then sonicated for 5 minutes with 30 second ON/OFF at 75% amplitude. Pre-validated antibody against DDX41 (Cell Signaling Technology, D3F1Z) was then pre-coupled to Anti-Rabbit IgG Dynabeads (ThermoFisher), added to lysate, and incubated overnight at 4°C. Prior to immunoprecipitation, 2% of the sample was taken as the paired input sample, with the remainder magnetically separated and washed with eCLIP high stringency wash buffers. Immunoprecipitation (IP) and input samples were cut from the membrane at the relative band size to 75kDa above. RNA adapter ligation, IP-western, reverse transcription, DNA adapter ligation, and PCR amplification were performed as previously described^31^.

After sequencing, samples were processed with Eclipsebio’s proprietary analysis pipeline (v1). UMIs were pruned from read sequences using umi_tools (v1.1.1). Next, 3’ adapters were trimmed from reads using cutadapt (v3.2). Reads were then mapped to a custom database of repetitive elements and rRNA sequences. All non-repeat mapped reads were mapped to the human genome (hg38) using STAR (v2.7.7a). PCR duplicates were removed using umi_tools (v1.1.1). Peaks were identified within eCLIP samples using the peak caller CLIPper (v2.0.1). For each peak, IP versus input fold enrichments and p-values were calculated by the Yates’ Chi-Square test, or Fisher Exact Test if the observed or expected read number was below 5. Peaks were annotated using transcript information from GENCODE v41 with the following priority hierarchy to define the final annotation of overlapping features: protein coding transcript (CDS, UTRs, intron), followed by non-coding transcripts (exon, intron). Differential binding analysis comparing experimental conditions was performed using DESeq2 (v1.34.0).

### Electrophoretic Mobility Shift Assay (EMSA)

DDX41 wildtype and R525H proteins were purified as described previously^32^. RNA binding was assessed using a Cy5-labelled RNA substrate derived from the Minx pre-mRNA, containing a polypyrimidine tract and the 3’-exon^33^. The protein samples were serially diluted with dilution buffer (20 mM HEPES, pH 7.9, 200 mM KCl, 10% glycerol) and incubated for 10 minutes at room temperature in the absence of ATP, or in the presence of 0.2 mM ATP. The samples were then mixed with RNA and incubated for 20 minutes at room temperature. The final sample contained 18 mM HEPES, pH 7.9, 130 mM KCl, 10 nM RNA, 0.005 % NP40, and 6.5 % glycerol. Complexes were resolved on an 8% pre-cast native gel (Novex). Quantification was performed using the iBright Analysis software, with local background subtraction. For each gel lane the fraction of species observed for single, double, and multimeric complexes is calculated relative to the sum of all complexes and the free unbound RNA. Error bars were obtained from at least two independent purifications and two technical replicates for each purification.

### Animals

All animal experiments were performed with approval by and in accordance with Fred Hutchinson Cancer Center (FHCC) Institutional Animal Care and Use Committees (IACUC) guidelines. Animal experiments were performed within the FHCC Comparative Medicine facility. To model the truncating germline *DDX41* mutations, a constitutive heterozygous knockout mouse model (*Ddx41*^KO/+^) was generated by introducing an indel in exon 1 of *Ddx41* by CRISPR. *Ddx41*^KO/+^ mice were crossed with *Ddx41*^Δ/+^ mice and offsprings were genotyped at weaning (2-3 weeks of age). A conditional *Ddx41*^R525H^ allele was generated using a Cre/loxP targeting strategy. A floxed cDNA cassette spanning exons 11–17 was inserted upstream of the corresponding endogenous exons, flanked by loxP sites. The copy of exon 15 downstream of this cassette was engineered to carry a single-nucleotide substitution (GCA→GTA), encoding the R525H missense mutation. In *Ddx41*^Mx^^1-R525H/+^ mice, pIpC treatment induced hematopoietic-restricted Cre expression and recombination. Recombination and mutation status were confirmed by Sanger sequencing of bone marrow DNA. *Ddx41*^Mx^^1-R525H/+^ mice were crossed with *Ddx41*^KO/+^ mice to generate *Ddx41* biallelic mutant mice on pIpC induction. After 3 rounds of pIpC, end point analysis was performed from peripheral blood, bone marrow and spleen. For RNA-sequencing, Lin^-^cKit^+^ bone marrow cells were sorted, and RNA extraction and library prep were performed as described above.

### Patient samples

Cryopreserved de-identified specimens from MDS and AML patients were obtained from the Fred Hutchinson Cancer Center/University of Washington Hematopoietic Diseases Repository (FHCC/UW-HDR) and from the University of Pennsylvania repository. As part of the FHCC/UW-HDR, patient specimens are collected and stored under the oversight of Fred Hutch Institution Review Office, and all participants provided written informed consent that adhered to the guidelines of 1975 Declaration of Helsinki. Diagnosis for all the patients was confirmed using established guidelines at the time of diagnosis.

## RESULTS

### Somatic *DDX41* mutations fail to rescue cell proliferation of DDX41-deficinet AML cells and are functionally distinct from germline *DDX41* mutations

To systematically evaluate the function of pathogenic/likely-pathogenic (P/LP) *DDX41* mutations, we designed a series of DDX41 cDNAs that were wildtype (WT), or that carried specific non-truncating germline mutations (G173R, E256K, Y259C, and S363del) or somatic mutations (R525H and G530D) (**Figure 1A**, top). We performed proliferation assays using genetic complementation in isogenic MOLM-13 AML cell lines engineered to express a doxycycline-inducible shRNA against the 3’ UTR of endogenous *DDX41*, which selectively depletes endogenous DDX41 transcript without targeting ectopic DDX41 cDNAs. Western blot analysis from MOLM-13 cells revealed a drastic reduction in DDX41 protein level 48 hours after doxycycline treatment (**Figure S1A**). DDX41-knockdown cells were complemented with empty vector (EV), DDX41 WT or mutant cDNAs (**Figure 1A**, bottom). In the absence of doxycycline, we did not observe any appreciable differences in growth (**Figure S1B and S1C**). After doxycycline induction, we found that knockdown of *DDX41* inhibited proliferation of MOLM-13 cells, which was not rescued by empty vector (**Figure 1B and 1C**). Re-expression of *DDX41-*WT cDNA fully rescued the proliferation defects in DDX41-knockdown cells (**Figure 1B**), restoring live cells numbers to levels comparable to uninduced cells (**Figure S1B**). Notably, overexpression of all four DDX41 germline mutant cDNAs also fully rescued proliferation in DDX41-knockdown cells to levels indistinguishable from cells expressing DDX41-WT cDNA (**Figure 1B**). In contrast to germline mutants, somatic *DDX41* hotspot mutants R525H and G530D failed to rescue proliferative defects in DDX41-knockdown MOLM-13 cells (**Figure 1C**).

**Figure 1:**
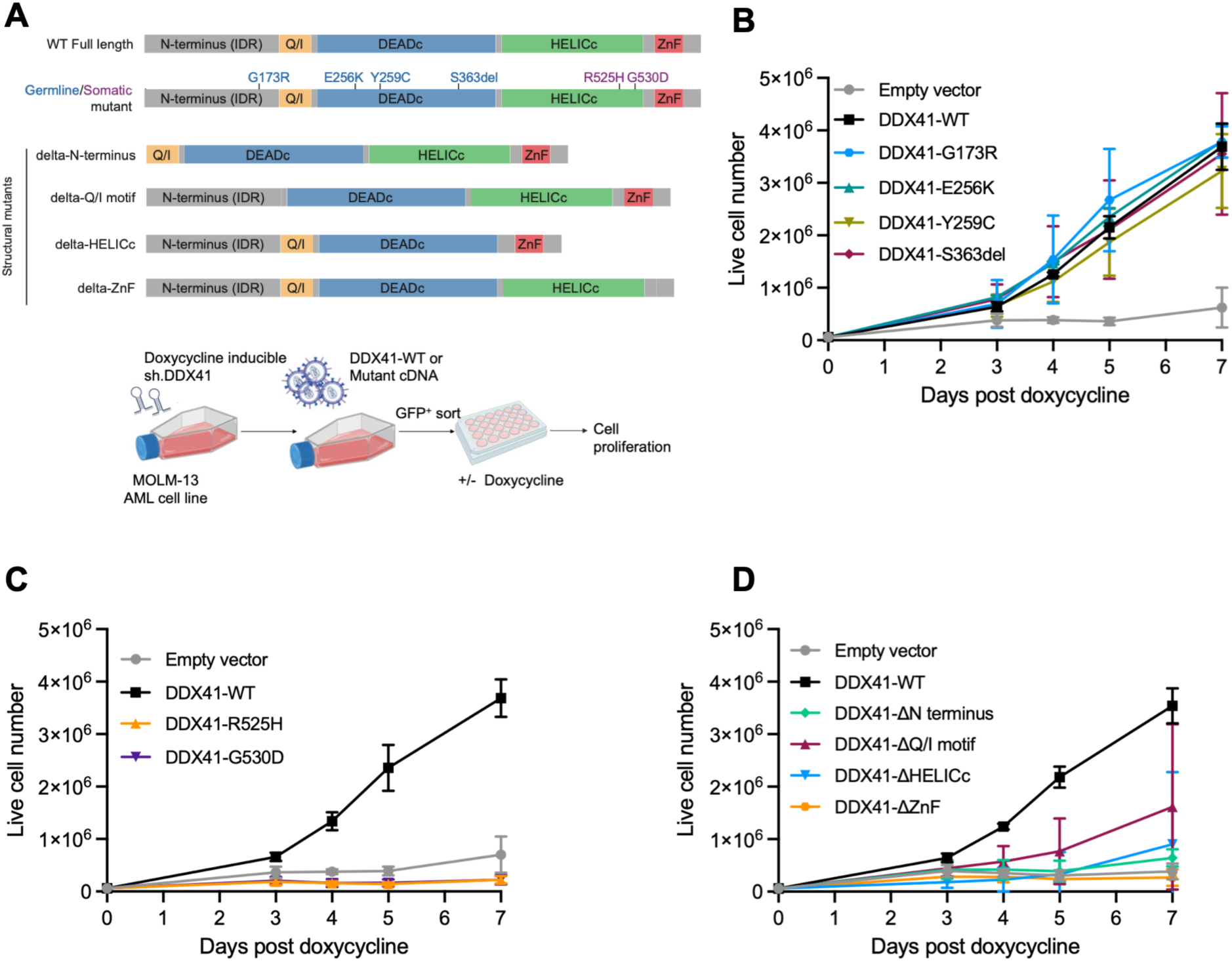
Somatic DDX41 mutations are functionally distinct from germline DDX41 mutations. **(A)** Schematic representation of DDX41 protein domains, pathogenic non-truncating germline or somatic mutations screened and DDX41-domain deletion mutants (top) and cell proliferation assay in MOLM-13 cells (bottom). Rescue of cell proliferation in presence of doxycycline by **(B)** germline DDX41 mutants, data are mean ± SD of n=3 independent experiments **(C)** somatic DDX41 mutants, data are mean ± SD of n=4 independent experiments and **(D)** structural DDX41 mutants, data are mean ± SD of n=3 independent experiments.

To identify the functional domains of DDX41 that are required for its proliferative function, we next generated a panel of *DDX41* structural mutants lacking the N-terminal domain, the Q/I motif, the HELICc domain and the zinc-finger domain (ZnF) and tested their ability to rescue proliferation following DDX41 knockdown (**Figure 1A**, top, and **Table S3**). All four DDX41 domain deletion mutants failed to rescue proliferation of MOLM-13 cells after doxycycline induction, with delta-ZnF mutant showing the most severe defect (**Figure 1D**), indicating that multiple domains of DDX41 are critical for its function. Cells not induced with doxycycline and overexpressing different *DDX41* structural mutants did not show any growth defects (**Figure S1D**). Taken together, these data show that somatic *DDX41* mutations harbor loss-of-function properties and are functionally distinct from *DDX41* germline mutations in a genetic complementation model. The failure to rescue cell growth by any of the domain-deletion mutants highlights the essentiality of DDX41 in maintaining cellular fitness of AML cells.

### AML-associated somatic *DDX41* mutations R525H and G530D drive widespread splicing alterations in key HSC regulators

The *DDX41*-R525H mutation has been described to have decreased ATPase and helicase activity^7^. Given the role of DDX41 in splicing and its interaction with the spliceosome C* complex^19,20,26^, we hypothesized that somatic hotspot mutations R525H and G530D, both located within the helicase domain of DDX41, hinder RNA processing and cause mis-splicing of specific transcripts that, in turn, promote dysplastic hematopoiesis. To identify key RNA targets aberrantly spliced by *DDX41* mutants, we performed RNA sequencing and splicing analysis on isogenic MOLM-13 cells induced with doxycycline to trigger knockdown of DDX41, and over-expressing somatic or germline *DDX41* mutants (**Figure 2A**). Differential splicing analysis identified that RNA mis-splicing is ten times more enriched in the *DDX41* somatic mutants (**Figure 2B** and **Figure S2A**), with a total of 2400 mis-spliced events each in DDX41-R525H and G530D, compared to *DDX41* germline mutants with only a few hundred (**Figure S2B**). We focused our analysis on alternative 3’ splice site usage (A3SS), alternative 5’ splice site usage (A5SS), retained intron (RI) and skipped exons (SE) events promoted in *DDX41* somatic mutants R525H and G530D compared to wildtype. Using a delta percent spliced in (delta-PSI) cut-off of > +0.2 and FDR of <0.01, we ranked significant splicing events based on their delta-PSI for *DDX41*-R525H, *DDX41*-G530D and all *DDX41-*germline mutants (**Figures 2C**, **2D** and **Figures S2C through S2F**, respectively). This revealed that isogenic AML cells with *DDX41*-R525H or *DDX41*-G530D mutation not only had increased number of splicing events, but also a higher magnitude of significant splicing events compared to *DDX41* germline mutants (**Figures 2C**, **2D** and **Figures S2B through S2F**). Next, we generated a list of high-confidence mis-spliced genes enriched in *DDX41* somatic mutants for validation by RT-PCR by following these criteria: (a) mis-spliced events should overlap between *DDX41*-R525H and *DDX41*-G530D, and be excluded from all *DDX41*-germline mutants, and (b) the events should be discernable by visual inspection of coverage plots in Integrated Genomics Viewer (IGV) (**Figure 2E**). We found enrichment of targets with previously described or potential roles in hematologic malignancy or DDX41 biology, including CLK3^21,34^, SEPTIN7^35,36^, YTHDF1^37,38^, DVL1^39,40^ and RPL32^14^ (**Figures 2C** and **2D**). Validation of some of the top mis-spliced targets by visual inspection of coverage tracks in IGV (**Figures 2F** and **S2G**) and RT-PCR with isoform-specific primers (**Figures 2G**) confirmed mis-splicing of SEPTIN7, CLK3 and DVL1 in *DDX41* somatic mutants R525H and G530D only, but not in any of the *DDX41* germline mutants, *DDX41* wildtype or empty vector and non-targeting controls. To assess the impact of *DDX41* somatic and germline mutants on global transcription, we performed principal component analysis (PCA) on RNA-seq gene expression data. *DDX41* somatic mutants R525H and G530D show transcriptional separation from *DDX41* germline mutants and *DDX41* wildtype (**Figure S2H**), paralleling the splicing-level distinction observed (**Figure 2B**). Unsupervised hierarchical clustering of the union set of differentially expressed genes (DEGs) across *DDX41* mutants further confirmed that *DDX41* somatic mutants R525H and G530D showed the strongest divergence from wildtype, while the *DDX41* germline mutants retained more similarity to wildtype (**Figure S2I**). To isolate the transcriptional changes common to both somatic mutants, we examined the intersection of genes differentially expressed in DDX41-R525H vs. wildtype and DDX41-G530D vs. wildtype. Clustering of this gene set revealed that gene expression from the two somatic mutants were highly concordant to each other and divergent from wildtype (**Figure S2J**) indicating that the *DDX41* somatic mutants drive broader transcriptional dysregulation beyond splicing alone. Taken together, these results indicate a splicing and transcriptional signature unique to *DDX41* somatic mutants, corroborating the functional differences observed between *DDX41* somatic and germline mutants in AML cells.

**Figure 2:**
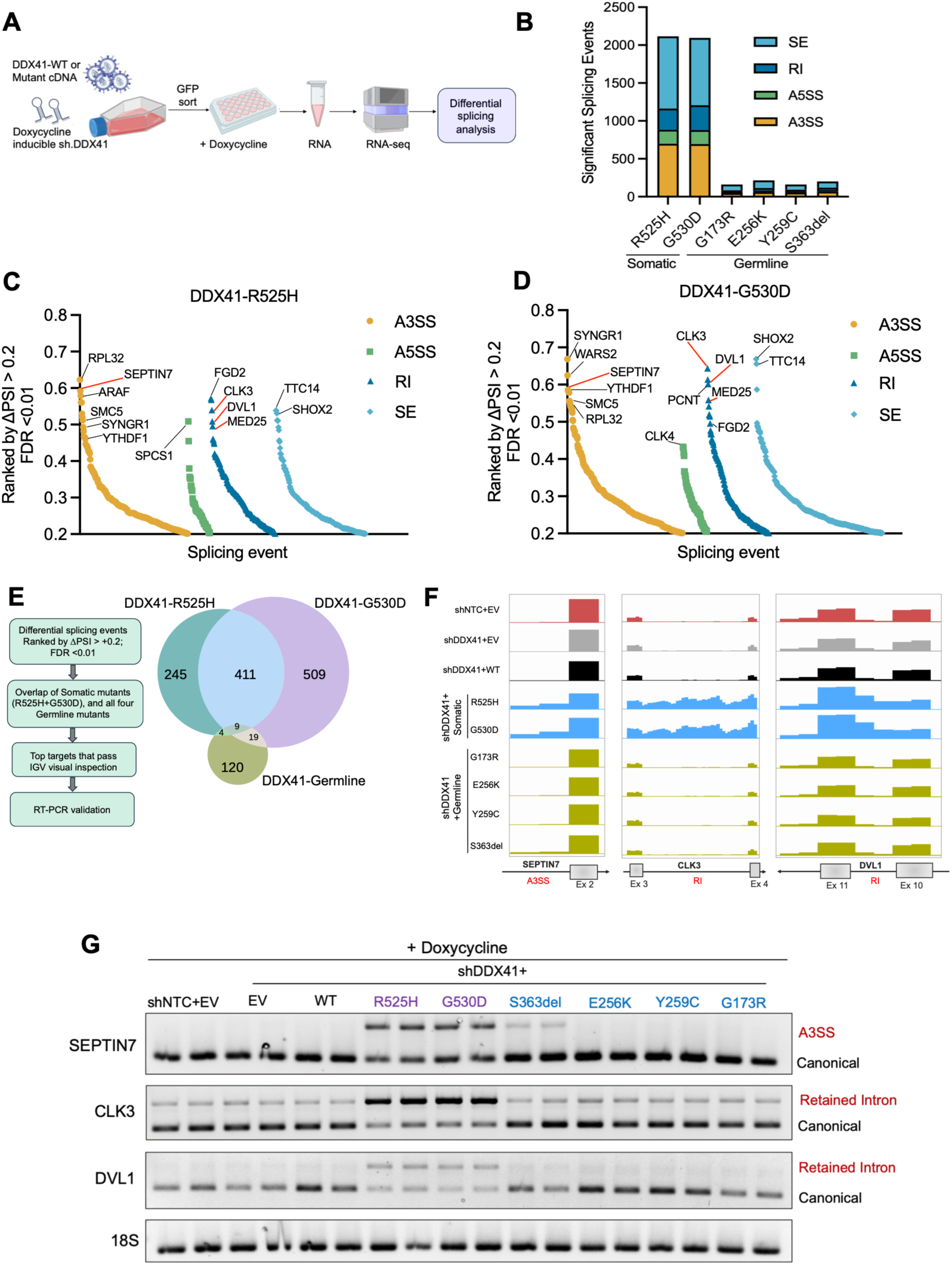
DDX41 somatic hotspot mutants R525H and G530D induce a distinct, high-magnitude mis-splicing signature relative to germline mutants. (A) Schema representing RNA-sequencing workflow in isogenic MOLM-13 cells. (B) Significant splicing events across different DDX41 mutants relative to DDX41 wildtype, filtered as FDR < 0.05 and |delta-PSI| > 0.2. PSI and FDR adjusted for multiple comparisons were calculated using rMATS. A3SS=alternative 3’ splice site usage, A5SS=alternative 5’ splice site usage, RI=retained intron, SE=skipped exon. Significant splicing events ranked by delta-PSI > 0.2 and FDR < 0.01, promoted in (C) DDX41-R525H and (D) DDX41-G530D. (E) Flow chart of criteria used to prioritize top targets for validation (left) and Venn diagram showing overlap of genes mis-spliced in DDX41 somatic and germline mutants (right). (F) IGV coverage plots of statistically significant mis-spliced events: alternative 3’ splice site usage of SEPTIN7, retained intron of CLK3 and DVL1. Y-axis, RPMK normalized coverage. (G) RT-PCR of select targets (SEPTIN7, CLK3, DVL1) using isoform specific-primers across DDX41 mutants, wildtype and empty vector controls. 18S, loading control.

### Mis-spliced genes are enriched among transcripts bound by DDX41 R525H

To determine whether the transcripts mis-spliced by DDX41 R525H are also direct RNA binding targets of mutant DDX41, we performed enhanced crosslinking immunoprecipitation (eCLIP) in MOLM-13 cells over-expressing DDX41-WT or DDX41-R525H cDNA (**Figure 3A**). Metagene analyses showed that DDX41 peaks, for both WT and R525H, were concentrated within coding exons (**Figure S3A**). We overlapped RNA-targets reproducibly bound across duplicate samples to the set of transcripts mis-spliced in DDX41-deficient MOLM-13 cells re-expressing DDX41 R525H (sh.DDX41+R525H) and found that of the 1,566 R525H-bound genes, 231 (14.8%) were also mis-spliced (**Figure 3B**). Comparison of this association with DDX41-WT bound genes revealed a set of 148 genes that were uniquely bound and mis-spliced by DDX41-R525H (**Figure 3C**). Inspection of RBP peaks on IGV confirmed an enriched binding peak in R525H immunoprecipitation (IP), absent in WT-IP or size-matched input controls for SEPTIN7 at exon 11 and MED25 at exon 15 (**Figure 3D**). Gene Ontology analysis of the 148 genes bound and mis-spliced by R525H revealed a significant enrichment of terms related to RNA splicing, chromatin remodeling and regulation of intracellular transport, among others (**Figure S3B**). These data illustrate that somatic DDX41-R525H binds RNA transcripts that are enriched among those mis-spliced by the mutant and critical to cellular homeostasis.

**Figure 3:**
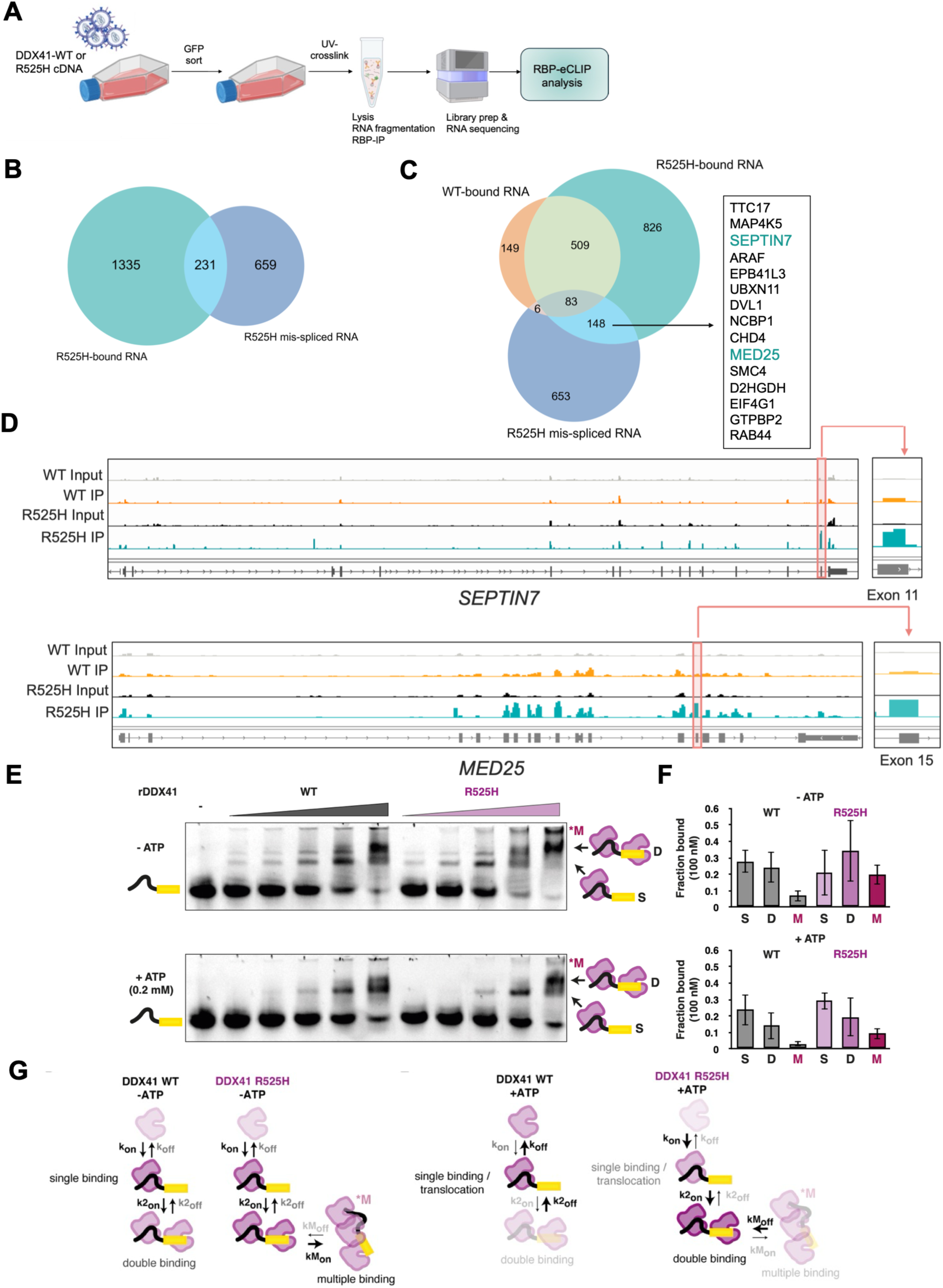
DDX41 R525H alters RNA binding on mis-spliced transcripts and shifts the mode of RNA engagement *in vitro*. **(A)** RBP-eCLIP workflow in MOLM-13 cells expressing DDX41-WT or R525H cDNA. **(B)** Overlap of R525H-bound and R525H mis-spliced transcripts. **(C)** Overlap of WT-bound, R525H-bound, and mis-spliced transcripts; R525H-specific bound and mis-spliced genes listed at right. **(D)** eCLIP coverage (input and IP) at SEPTIN7 and MED25, with insets at exon 11 and exon 15 (red boxes). **(E)** EMSAs with increasing recombinant DDX41 WT or R525H, ±0.2 mM ATP; single (S), double (D), and multiply bound (*M) complexes indicated. **(F)** Fraction bound in each state at 100 nM protein. **(G)** Model of binding transitions for WT and R525H ±ATP; arrows indicate relative changes in rate constants.

DDX41 is predicted to bind RNA within a conserved cleft formed by the DEAD and HELICc domains, with ATP binding stabilizing the closed RNA-bound conformation (**Figure S3C**, left), as observed for other DEAD-box ATPases^25,41–43^. Additionally, the R525H hotspot mutation localizes within motif VI adjacent to the ATP-binding pocket (**Figure S3C**, right). To understand how DDX41 engages physiologically relevant RNA sequences, we analyzed its binding to an RNA containing the Minx pre-mRNA polypyrimidine (PY) tract, 3′-splice site (3′-ss) and 3′-exon. EMSA experiments *in vitro* showed that WT DDX41 binds RNA as discrete single-(S) and double-bound (D) complexes, with a minor population of slower-migrating higher-order (M) assemblies (**Figure 3E**), which may represent multimeric filaments or aggregates. Addition of ATP substantially reduced formation of higher-order assemblies, consistent with ATP-driven conformational cycling causing an increase in the dissociation rate (k_off_) and promoting release and remodeling of RNA-bound complexes, as seen in other typical DEAD-box ATPses^42,43^. Across multiple independent experiments, our data suggest that increasing DDX41 concentration causes redistribution between multiple RNA-bound states. Thus, although the apparent Kd is in the 100-200 nM range, fitting the EMSA data to a single equilibrium dissociation constant provided limited mechanistic insights since increasing concentrations cause a change in relative partitioning of S, D, and M species. We chose instead to report the fraction of specific RNA-bound complexes at a given limiting concentration of 100 nM (**Figures 3E** and **3F**). These observations suggest that DDX41 normally undergoes rapid ATP-dependent cycles of RNA binding and release. Such cycling is likely to be essential for DDX41 function, either by remodeling structured RNA substrates such as R-loops or by promoting dynamic rearrangements during spliceosome assembly and catalysis. We propose that ATP hydrolysis normally increases the effective dissociation rate of DDX41, preventing prolonged accumulation of multi-protein RNA-bound assemblies (**Figure 3G**). The R525H mutation instead slows ATP-dependent release, kinetically trapping mutant DDX41 on RNA and allowing progressive accumulation of higher-order complexes. Therefore, the R525H mutant potentially converts a transient RNA-remodeling enzyme into a persistent RNA-bound scaffold and compromises accurate splicing, potentially by disrupting spliceosome dynamics.

### *DDX41*-R525H and *DDX41*-G530D mutations confer neomorphic functions

Next, we questioned if *DDX41-R525H* and *DDX41-G530D* reflected loss or gain of function. We used alanine mutagenesis to generate R525A and G530A mutants and assayed these mutants for cell growth in the same isogenic system described in Figure 1. Inhibition of cell growth triggered by doxycycline induced knockdown of DDX41 was not rescued by the alanine mutants R525A and G530A, with R525A behaving similarly to empty vector rescue and G530A showing partial rescue of growth at day 7 (**Figure 4A**), indicating a loss-of-function phenotype. Cells not induced with doxycycline and overexpressing alanine substitution mutants did not show any growth defects (**Figure S4A)**. Strikingly, RT-PCR of validated mis-spliced targets SEPTIN7, CLK3 and DVL1 revealed that R525A and G530A substitutions did not induce aberrant splicing (SEPTIN7, DVL1) or perturbed splicing to lower levels (CLK3) than R525H and G530D (**Figure 4B**). Thus, aberrant splicing is a gain-of-function conferred specifically by the disease-associated mutations at residues R525 and G530.

**Figure 4:**
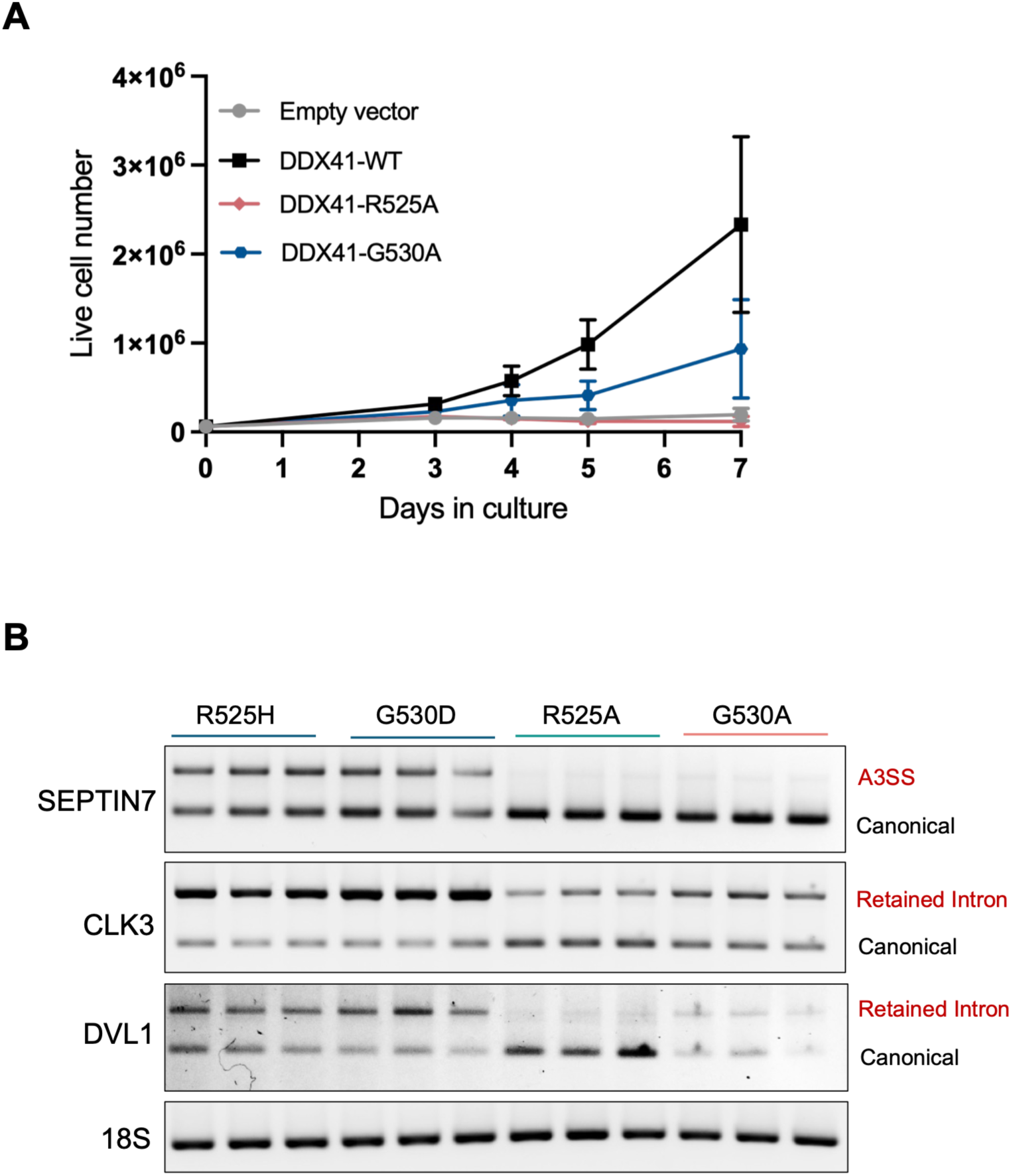
DDX41-R525H and DDX41-G530D mutations confer neomorphic functions. **(A)** Rescue of cell proliferation in presence of doxycycline by alanine mutants R525A and G530A, data are mean ± SD of n=3 independent experiments. **(B)** RT-PCR of SEPTIN7, CLK3 and DVL1 in cells expressing empty vector, R525H, G530D, R525A, or G530A, showing canonical products and the mutant-associated mis-spliced isoforms. 18S, loading control.

### *Ddx41*^KO/R525H^ mice show severe bone marrow failure, and the biallelic mutation induces global splicing changes in mice

We next sought to verify the effects of the somatic *DDX41* R525H mutation in a physiologically relevant context that faithfully captures the genetic configurations seen in patients. To mimic *DDX41* frameshift truncating mutations (e.g. *DDX41*-D140fs and *DDX41*-A500fs) that account for >60% of germline variants, we first generated a constitutive heterozygous knockout mouse model (*Ddx41*^KO/+^) by introducing an indel in exon 1 of *Ddx41* by CRISPR (**Figure S5A**). To determine whether our germline *Ddx41* mutant allele behaved as a null, we intercrossed *Ddx41*^KO/+^ mice and genotyped offspring at weaning (2-3 weeks of age). We did not recover *Ddx41*^KO/KO^ animals from n= 30-40 pups (expected: n=∼10; χ^2^ p= 0.0006), indicating the homozygosity of our mutant allele is incompatible with survival (**Figure S5A**). This is consistent with a previous report in which germline deletion of *Ddx41* failed to yield homozygous null pups^44^.

Next, to model *DDX41* somatic mutations (e.g. *DDX41-*R525H) that are acquired later in disease pathogenesis exclusively in hematopoietic cells, we generated a novel conditional *Ddx41*-R525H knock-in model that expresses the somatic variant from the endogenous murine *Ddx41* locus (**Figure S5B**). Briefly, a minigene containing *Ddx41* exons 11-17 flanked by two loxP sites was inserted into *Ddx41* intron 10 to allow for expression of wildtype *Ddx41*. In addition, the R525H point mutation (C**<u>G</u>**T-->C**<u>A</u>**T) was introduced into exon 15 of endogenous *Ddx41*, which will be expressed upon Cre-mediated recombination and deletion of the minigene. We next crossed heterozygous *Ddx41*^R525H/+^ mouse with the *Mx1*-cre transgenic mouse strain^45^ to generate *Mx1*-Cre^Tg/+^ *Ddx41*^R525H/+^ mice (herein denoted as *Ddx41*^Mx^^1-R525H/+^), to induce expression of Cre recombinase specifically in hematopoietic stem and progenitor cells (HSPCs) by polyinosine-polycytosine (pIpC) administration as described previously^46^. Efficient recombination and expression of the *Ddx41*^R525H^ allele was confirmed by Sanger sequencing of Ddx41 cDNA extracted from bone marrow mononuclear cells (BM MNCs) 7-10 days post pIpC administration (**Figure S5B**).

We next crossed *Ddx41*^KO/+^ and *Ddx41*^Mx^^1-R525H/+^ mice to generate biallelic *Ddx41*-mutant (*Ddx41*^KO/Mx^^1-R5252H^) mice to mimic the genetic configurations found in human *DDX41*-mutant patients. Three days after the last dose of pIpC administration, we harvested bone marrow, peripheral blood and spleen for endpoint phenotyping (**Figure 5A**). Peripheral blood analysis from *Ddx41*^KO/Mx1-R525H^ biallelic mutant mice developed significant leukopenia (**Figure 5B)** and thrombocytopenia (**Figure 5C**), while neither heterozygous germline (*Ddx41*^KO/+^) nor the somatic R525H allele alone (*Ddx41*^Mx1-R525H/+^) showed any perturbations in white blood cell and platelet counts relative to *Ddx41* wildtype (*Ddx41*^+/+^) mice. Red blood cell counts remained unchanged across all four genotypes (**Figure 5D**). We observed severe bone marrow hypocellularity (**Figure 5E**) and depletion of hematopoietic stem and progenitor cells (defined as Lin^-^cKit^+^) (**Figure 5F**) in *Ddx41*^KO/Mx^^1-R525H^ mice relative to *Ddx41*^+/+^, *Ddx41*^KO/+^, or *Ddx41*^Mx^^1-R525H/+^ mice. Our data is consistent with prior reports of impaired hematopoiesis in *Ddx41*-deficient mice^14^, with the added distinction of true germline deletion of *Ddx41* in our model.

**Figure 5:**
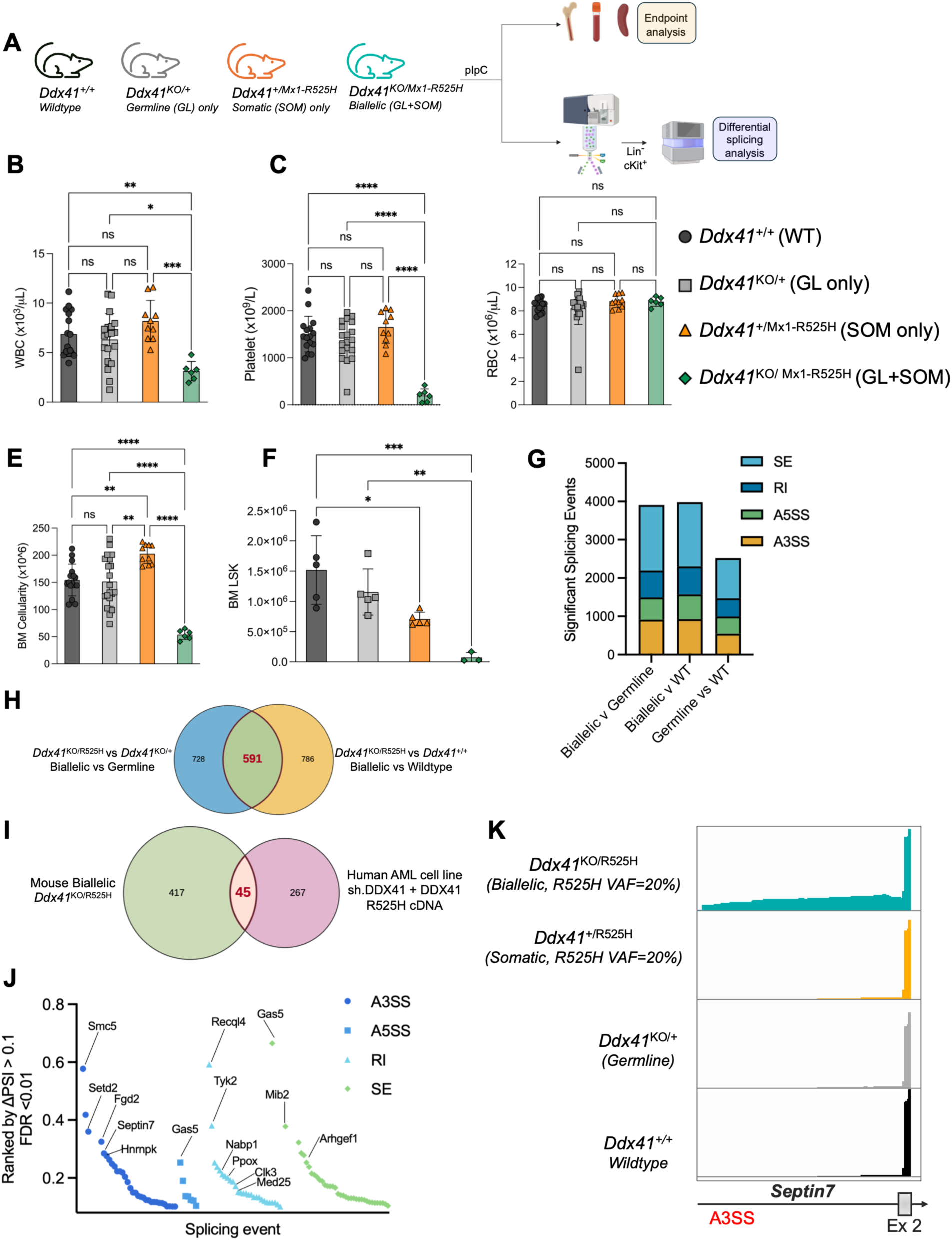
Biallelic *Ddx41*^KO/R525H^ mutants cause rapid and severe bone marrow failure, and the R525H splicing program is partially conserved between mice and humans. (**A**) Schematic of mouse model indicating genotypes of experimental groups followed by end point analysis and RNA-sequencing post pIpC. (**B–D**) Peripheral blood white blood cell, platelet, and red blood cell counts by genotype. (**E, F**) Bone marrow cellularity and LSK numbers. Bars show mean ± SD; ns, not significant; *p<0.05, **p<0.01, ***p<0.001, ****p<0.0001. (**G**) Differential splicing analysis on Lin^-^cKit^+^ sorted bone marrow showing numbers of significant splicing events by type (SE, RI, A5SS, A3SS) for each comparison, filtered as FDR < 0.05 and |delta-PSI| > 0.1. PSI and FDR adjusted for multiple comparisons were calculated using rMATS. (**H)** Overlap of mis-spliced genes enriched in biallelic mice relative to germline and to WT; 591 genes overlap. **(I)** Overlap of mis-spliced genes between biallelic mice and human DDX41 R525H isogenic MOLM-13 cells; 45 genes from the 591 set overlap. **(J)** Splicing events from the 45 species-conserved genes in biallelic mice ranked by delta-PSI > 0.1, FDR < 0.01. **(K)** IGV coverage tracks at *Septin7* showing the A3SS included in HSPCs of biallelic but not somatic only, germline only or WT mice.

Given our observation of global splicing perturbation by “biallelic” *DDX41-R525H* (germline-null captured by shRNA mediated *DDX41* knockdown + *R525H* configuration) in human AML cells, we asked if impaired hematopoiesis observed in *Ddx41*^KO/Mx^^1-R525H^ biallelic-mutant mice is accompanied by a distinct splicing signature. To this end, we sorted Lin^-^cKit^+^ HSPCs from pIpC induced mice and performed RNA-sequencing and differential splicing analysis (**Figure 5A**). We focused on the clinically predominant and disease-related biallelic representation of *DDX41* and quantified differential splicing in *Ddx41*^KO/Mx^^1-R525H^ biallelic-mutant mice relative to their wildtype (*Ddx41^+/+^*) and heterozygous germline (*Ddx41*^KO/+^) counterparts (**Figure 5A**). Indeed, both biallelic contrasts – biallelic versus germline and biallelic versus wildtype – yielded comparable and substantial numbers of significantly spliced events (FDR<0.01, |delta| PSI > 0.1), exceeding the germline-versus wildtype contrast, similar to our earlier findings in human AML cells (**Figure 5G**). To identify mis-splicing events promoted in the biallelic state independent of the comparator, we intersected the biallelic-versus-germline and biallelic-versus-wildtype event sets (FDR<0.01, ΔPSI > +0.1), defining a core set of 591 high-confidence shared mis-spliced genes in *Ddx41*^KO/MX^^1-R525H^ biallelic mutant mice (**Figure 5H**). GO analysis of the 591-core against the rMATS-testable background revealed enrichment of genes involved in genome maintenance (mismatch repair, DNA damage response), RNA processing (nonsense-mediated decay, mRNA deadenylation, and splicing regulation), and stem/progenitor programs (stem cell proliferation, β-catenin signaling) (**Figure S5C**), linking the mis-splicing program from biallelic mutant cells to pathways central to hematopoietic stem cell function. Cross-species comparison of this 591-core biallelic program against differentially spliced genes gained in our biallelic DDX41-*R525H*-specific dataset in human AML cells identified 45 conserved targets (14.4% of human genes), indicating a mis-splicing phenotype partially preserved between mouse and human (**Figure 5I**). To determine if our top validated mis-spliced targets from *DDX41-*R525H expressing human AML cells were recapitulated in mice, we ranked the 45-core genes by event type and effect size (ΔPSI > 0.1, FDR < 0.01) and examined the identity of the strongest events. Indeed, the 45-core recapitulated the top validated targets in our human cell line dataset (*Septin7, Clk3, Fgd2, Med25*), albeit at lower rank (**Figure 5J**). Among the most strongly mis-spliced events in mice are regulators of hematopoiesis (*Recql4*^47^), hematopoietic stem cell self-renewal (*Setd2*^48,49^), aging (*Smc5*^50^), and proliferation (*Hnrnpk*^51^) (**Figure 5J**). To validate rMATS-predicted events orthogonally, we inspected read coverage of the top-ranked 45-core targets using IGV. Of these, *Septin7* showed unambiguous biallelic mutant mice-specific inclusion of the predicted A3SS event, (not seen in *Ddx41*^Mx1-R525H/+^*, Ddx41*^KO/+^ or *Ddx41*^+/+^ mice; **Figure 5K**), while *Clk3* showed increased intron retention in biallelic mutant cells but with less distinct genotype separation (**Figure S5D**). Further, *Septin7* and *Clk3* mis-splicing events were conserved with the human locus. *Setd2* showed biallelic mutant mice-specific coverage consistent with the predicted A3SS event (**Figure 5D**), although the mis-splicing event was not conserved with human data. Taken together, our data show that biallelic *DDX41* mutations drive a distinct mis-splicing program, partially conserved between human and mouse and enriched for pathways such as genome maintenance, RNA processing, and hematopoietic stem cell pathways, linking splicing dysregulation to impaired hematopoiesis in the biallelic state.

### SEPTIN7 mis-splicing is enriched in CD34⁺ stem and progenitor cells of biallelic DDX41-mutant MDS/AML

To determine whether the mis-splicing events identified in our *DDX41*-mutant isogenic AML cell lines and mouse models also occur in patients, we collected primary MDS/AML bone marrow aspirates from three patient populations: a) patients without a *DDX41* mutation (wildtype), b) patients with a germline *DDX41* mutation only (GL only), and c) patients with biallelic *DDX41* mutations (GL+R525H) (**Figures 6A and S6A**). Only treatment-naïve patients were included, and assignment to the GL-only or GL+R525H group was based on clinically annotated next-generation sequencing data and clinically reported variant allele frequencies (VAFs). For each patient we isolated CD235a^-^CD45⁺CD34⁺ hematopoietic stem and progenitor cells (HSPCs) from marrow aspirates and validated the top mis-spliced targets identified in human DDX41-R525H AML cells by RT-PCR. In these cells, we observed the SEPTIN7 mis-splicing product (the A3SS isoform) restricted to GL+R525H (biallelic) cells, but not in GL only or wildtype samples (**Figure 6B**). PCR product corresponding to the retained intron event in DVL1 was present in biallelic mutant samples but was also observed at lower levels in GL only or wildtype samples. Likewise, CLK3 mis-splicing was not restricted to a genotype but was present across all samples (**Figure 6B**). Strikingly, the SEPTIN7 A3SS event was confined to the HSPC compartment and was absent in the paired CD235a^-^CD45⁺CD34^-^mature cell fraction across patients spanning a wide range of R525H VAFs (3-28%) (**Figure 6C**), indicating that the SEPTIN7 mis-splicing is a feature of the disease relevant stem and progenitor population. Based on these findings we identified SEPTIN7 as a unique target mis-spliced by the DDX41-R525H biallelic mutation status, conserved across humans and mice. Further, the 3’ intron sequence harboring the 144 bp inclusion arising from the A3SS event is highly conserved between human and mice (**Figure S6B**), revealing that SEPTIN7 A3SS mis-splicing is an evolutionarily conserved consequence of DDX41-R525H dysfunction rather than a species-specific event.

**Figure 6:**
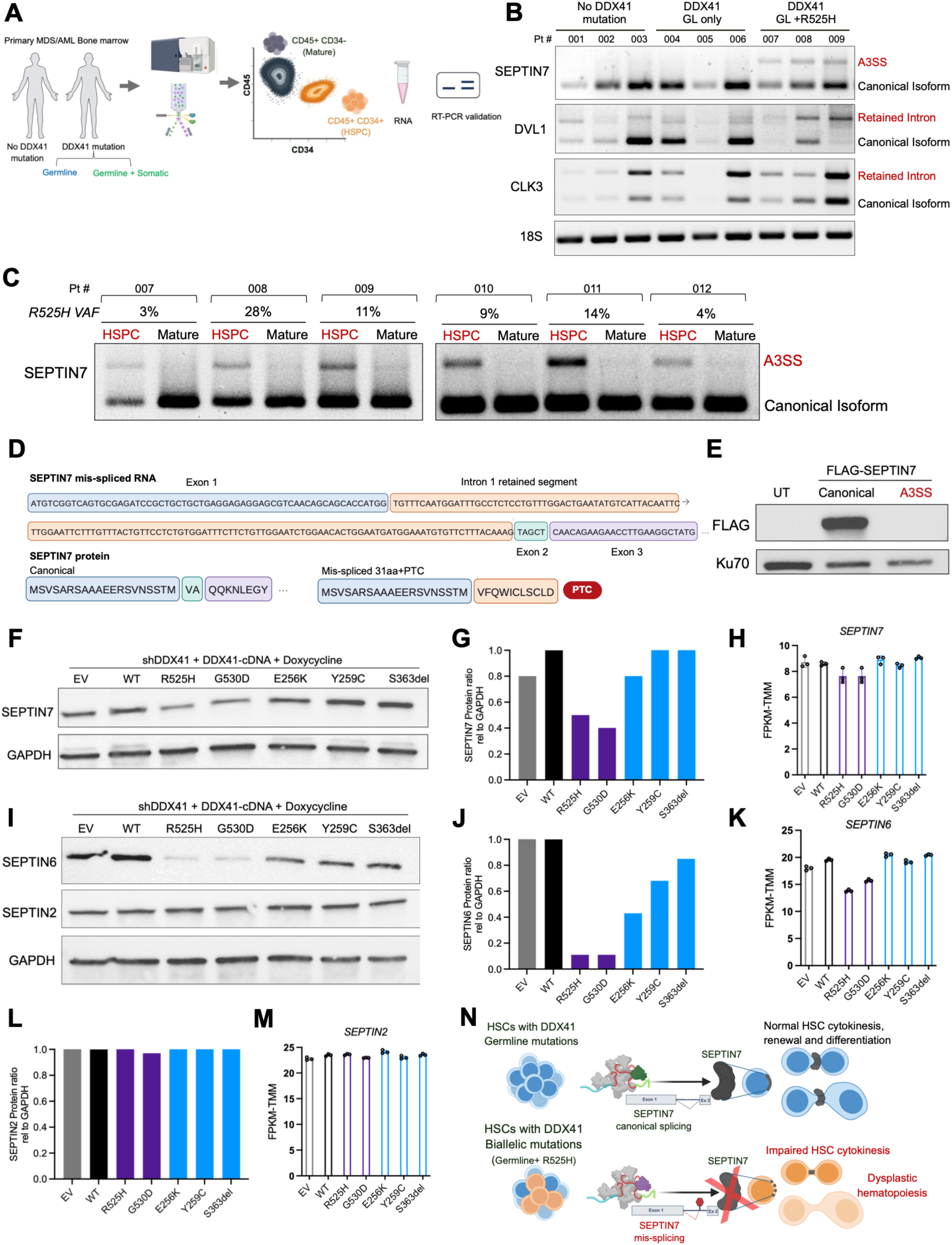
SEPTIN7 mis-splicing is enriched in CD34⁺ stem and progenitor cells of biallelic *DDX41*-mutant MDS/AML and leads to loss of SEPTIN7 protein. (**A**) Schematic of primary MDS/AML bone marrow harvest and sorting of CD45^+^ CD34^+^ HSPCs and CD45^+^ CD34^-^mature cells. (**B**) RT-PCR of SEPTIN7, DVL1, and CLK3 across patient groups, showing canonical products and the mutant-associated mis-spliced isoforms. 18S, loading control. (**C**) SEPTIN7 RT-PCR in paired HSPC and mature fractions from biallelic (GL+R525H) *DDX41* mutant patients with the indicated R525H VAF. (**D**) Schema of SEPTIN7 mis-spliced RNA and subsequent protein translation introducing a premature termination codon (PTC). (**E**) Western blot of untransduced (UT) cells and cells expressing FLAG-tagged canonical or A3SS SEPTIN7, probed for FLAG; Ku70, loading control. (**F–H**) SEPTIN7 western blot, quantification relative to GAPDH, and SEPTIN7 transcript levels (FPKM-TMM) in DDX41-knockdown cells re-expressing the indicated DDX41 cDNAs. (**I–K**) and (I, L, M) As in F–H for SEPTIN6 and SEPTIN2 respectively. (**N**) Model: in germline-only DDX41 mutation, canonical SEPTIN7 splicing supports normal HSC division and differentiation, whereas biallelic mutation drives SEPTIN7 mis-splicing, impaired HSC cytokinesis and accumulation of dysplastic HSCs.

We next asked how the A3SS event alters SEPTIN7. The inclusion of the 144 bp of the 3’ proximal intronic sequence is predicted to introduce a premature termination codon (PTC) (**Figure 6D**). Consistent with this, a FLAG-tagged construct containing the SEPTIN7-A3SS-specific isoform did not produce detectable protein, whereas the construct containing the canonical SEPTIN7 isoform was readily expressed (**Figure 6E**), establishing that the A3SS isoform does not generate a stable SEPTIN7 protein. To test whether this translates into loss of SEPTIN7 in a controlled genetic background, we used isogenic AML cells with an inducible shRNA targeting DDX41 and a series of DDX41-cDNAs expressing wildtype or mutant DDX41. SEPTIN7 protein was specifically reduced by ∼50% by the somatic (R525H and G530D) alleles, but not by the germline mutants E256K, Y259C, or S363del (**Figure 6F, 6G**). This reduction was largely post-transcriptional as SEPTIN7 mRNA levels were maintained across all alleles (**Figure 6H**), consistent with diversion of SEPTIN7 to the non-productive A3SS isoform rather than transcriptional downregulation. Notably, alanine mutants R525A and G530A did not result in the loss of SEPTIN7 protein in our genetic complementation model (**Figure S6C and S6D**), further emphasizing that the disease associated mutations R525H and G530D have neomorphic function.

SEPTIN7 is the central, non-redundant subunit of the Septin core complex and is required for assembly of the septin hetero-hexameric and hetero-octameric filaments that form the characteristic Septin ring essential for the completion of cytokinesis^52^. SEPTIN7 has also been reported to play a role in maintaining HSC polarity by regulating distribution of polarity proteins CDC42 and tubulin, and loss of *Septin7* in mice leads to failure of bone marrow repopulation of mice after lethal irradiation^35^. Because core septins co-stabilize each other and loss of one can destabilize its partners at the protein level^53^, we asked whether biallelic R525H-driven SEPTIN7 loss affected its partner subunits. In R525H-and G530D-expressing cells, SEPTIN6 protein was markedly reduced (**Figure 6I, 6J**), and although SEPTIN6 mRNA was also modestly decreased (**Figure 6K**), the protein loss appeared greater. SEPTIN2 protein and mRNA levels were unchanged across all alleles (**Figure 6I, 6L, 6M**). While preliminary, these observations are consistent with a selective effect on the SEPTIN7-SEPTIN6 module rather than a global change in septin expression and will require further study to establish whether SEPTIN6 is destabilized as a direct consequence of SEPTIN7 loss.

Taken together, biallelic DDX41-R525H drives HSPC-restricted mis-splicing of SEPTIN7 into a non-productive isoform and depletion of SEPTIN7 protein. Given the requirement for an intact septin complex in cytokinesis and stem-cell division, these findings suggest that SEPTIN7 loss may disrupt septin-dependent hematopoietic stem-cell division and differentiation (**Figure 6N**).

## DISCUSSION

The biallelic mutational state represents the defining genetic feature of *DDX41*-associated myeloid neoplasms. A germline lesion typically constitutes the first allelic event and the acquisition of a somatic mutation in *trans* – most often at the R525H hotspot in the second allele – indicates progression to overt disease. Whether these two classes of alleles are functionally distinct or simply represent different degrees of DDX41 loss, has not been delineated, and therefore requires a direct comparison of germline and somatic allele function. We use a genetic complementation strategy in isogenic human AML cells, in which endogenous DDX41 is silenced by a 3’ UTR-targeting shRNA and the function is restored by re-expression of P/LP *DDX41* mutant cDNA. We established that non-truncating germline and somatic *DDX41* mutations are functionally distinct. All four germline *DDX41* missense mutants tested (G173R, E256K, Y259C and S363del) rescued the proliferative defect of DDX41-depleted AML cells, indicating that these alleles retain proliferative function in isolation, whereas the somatic variants R525H and G530D failed to rescue proliferation defects in AML cells. This is consistent with the clinical observation that *DDX41* germline carriers sustain largely normal hematopoiesis for decades and provides a functional basis for why a second somatic hit is generally required for leukemic transformation. A recent preprint reached the same conclusion, showing that germline and somatic *DDX41* mutations are not functionally equivalent using a murine leukemia model^54^. Structure-function analysis of DDX41 domain-deletion mutants showed failure of proliferative rescue for all mutants, which is not unexpected for an essential gene. Notably, the R525 and G530 residues lie immediately upstream of the C-terminal zinc-finger (ZnF) domain – which is unique to DDX41 among DEAD-box helicases – and is predicted to cap RNA binding channel (**Figure S3C**). This raises the possibility that *DDX41* somatic mutations disrupt a C-terminal function required for RNA engagement, a hypothesis that will require further testing.

DDX41 is a component of the catalytically active spliceosome^19,26^ and coordinates pre-mRNA splicing^3,17,20^. To define the mechanism that distinguishes germline and somatic *DDX41* alleles, we profiled global splicing of DDX41-depleted AML cells reconstituted with wildtype or mutant DDX41 cDNA. Differential splicing analysis revealed that the somatic mutants R525H and G530D each induced an order of magnitude more significant splicing changes than any of the germline variants tested. Notably, both somatic mutants converge on a shared set of mis-splicing events, arguing for a common mechanism rather than allele-specific effects, and these events were enriched for A3SS usage together with intron retention, a signature consistent with the proposed roles of DDX41 in 3′ splice site selection^19,20^. Stringent filtering defined a somatic-specific splicing signature comprising established regulators of hematopoiesis (e.g. CLK3, SEPTIN7, YTHDF1, and RPL32), alongside genes with plausible HSC-related roles (e.g. DVL1 and MED25). We also provided evidence in support of altered RNA engagement by DDX41 somatic mutant protein rather than a downstream consequence of impaired proliferation. First, analysis showed that mis-spliced RNAs were significantly enriched among transcripts bound by DDX41-R525H but not by transcripts bound exclusively by wildtype DDX41, indicating that aberrant splicing could in part be a direct consequence of altered binding of RNA by DDX41 somatic mutant proteins. Notably, several of our top genes validated for mis-splicing like SEPTIN7 and MED25 showed increased binding by the R525H mutant compared to wildtype. Second, recombinant DDX41-R525H bound RNA aberrantly, accumulating higher-order complexes that, unlike wildtype DDX41, failed to resolve these complexes in the presence of ATP. DEAD-box helicases remodel RNA-protein complexes in an ATP-dependent manner^55,56^, and for several DEAD-box helicases, ATP binding without subsequent hydrolysis promotes stable RNA binding with half-lives of many hours, a process known as RNA clamping^55,56^. We therefore propose a model where the ATPase-deficient DDX41-R525H acts as an RNA clamp, increasing binding and delaying release of RNA substrate and thereby deregulating spliceosome assembly and disassembly on bound RNA. Our splicing-centric model offers a distinct mechanistic explanation from prior work, which have focused on R-loop accumulation and increased inflammatory signaling^13^, dysregulated ribosome biogenesis, snoRNA processing and translational defects^7,14,15^, and co-transcriptional RNA processing defects^17^. These mechanisms are not mutually exclusive – a defective RNA helicase with prolonged occupancy on nascent RNA could plausibly contribute to any of these mechanisms, whereas our data point to a defined somatic, allele-specific defect in pre-mRNA splicing.

A key question is whether the anti-proliferative and mis-splicing phenotypes of R525H and G530D variants reflect loss of function of the wildtype residue or is a specific property of the disease-associated substitutions. Alanine substation was used to separate these two phenotypes. DDX41-R525A failed to rescue proliferation of DDX41-depleted AML cells, comparable to R525H; whereas G530A partially rescued cell growth on day 7, suggesting that the R525H substitution is more severe in its role in cell proliferation. At the level of splicing, however, neither alanine mutant recapitulated the disease-associated phenotype: the aberrant SEPTIN7 (A3SS), CLK3 (RI) and DVL1 (RI) isoforms were strongly induced by the somatic mutants R525H and G530D, but only weakly, if at all, by R525A and G530A. These data argue that R525H and G530D are not simple loss-of-function alleles, as they are commonly interpreted, but confer a neomorphic function specific to splicing rather than a general gain-of-function, since it does not extend to the proliferative phenotype. How a histidine or aspartate substitution at these positions leads to aberrant splicing activity remains to be determined and is an important question for future work. It is worth noting that *DDX41* germline mutants also perturbed basal splicing in our system, albeit at far lower magnitude than somatic mutants. We speculate that this low-grade mis-splicing primes the hematopoietic environment over time, favoring selection of HSCs that acquire a HELICc-domain mutation in *trans* (e.g. R525H or G530D).

Individuals with germline *DDX41* mutations carry the mutant allele in every cell, and leukemic transformation is thought to require acquisition of a somatic mutation (most often R525H) in the second allele in hematopoietic cells^3,7^. Our novel *Ddx41* biallelic mutant mouse model faithfully mimics the genetic configuration found in patients – biallelic mutant mice develop bone marrow failure, recapitulating phenotypes reported previously using a model in which both germline and somatic mutations were confined to the hematopoietic compartment^14^. This indicates that bone marrow failure is a reproducible consequence of the *DDX41* biallelic state rather than a particular targeting strategy. The germline component of our strategy makes it possible to ask how a germline-mutant microenvironment shapes HSC behavior and second-hit selection, a question not readily addressable using hematopoietic-restricted models. Differential splicing analysis in physiologically relevant HSPCs revealed widespread mis-splicing in ‘germline + somatic’ *Ddx41* biallelic mutants relative to *Ddx41* germline-only mutants or *Ddx41* wildtype mice. This program was partially conserved with that of DDX41-depleted human AML cells re-expressing the R525H variant. Mis-spliced transcripts specific to the *DDX41* biallelic state (*Ddx41*^KO/Mx1-R525H^ in mice or sh.DDX41*+*DDX41-R525H in AML cells) across species included *bona fide* hematopoietic regulators (*Setd2, Septin7, Clk3, Recql4, Smc5* and *Hnrnpk*) and genes with plausible roles in hematopoiesis (*Fgd2* and *Med25*). Careful evaluation revealed that not all mis-spliced events were locus-conserved across species: examples-*Setd2* and *Recql4*. Whereas for *Septin7* and *Clk3,* mis-splicing was locus-conserved. This is consistent with the clamping model, in which prolonged occupancy by DDX41-R525H renders a bound transcript susceptible to mis-splicing.

A central limitation of engineered cell lines and mouse models is that they cannot, on their own, establish that a molecular event occurs in human disease. *DDX41*-mutant or wildtype bone marrow specimens from patients with MDS/AML stratifies natural genetics between germline carriers with and without the second somatic hit. RT-PCR on CD34^+^ HSPCs detected aberrantly spliced A3SS isoform of SEPTIN7 only in patients carrying a somatic R525H allele (GL+R525H), but not in patients with the germline lesion alone or with the wildtype *DDX41* allele. In contrast, the aberrantly spliced isoforms of CLK3 and DVL1 were not restricted to *DDX41* genotype, indicating mis-splicing of these transcripts are broader features of MDS and AML. This confirmed two things: a) the mis-spliced events identified and validated in our cell line model are not artifacts of cell engineering, but true events present in MDS/AML marrow, and b) SEPTIN7 mis-splicing is dependent on the second somatic hit, marking it as a *DDX41*-biallelic (GL+R525H) genotype specific readout, rather than an MDS/AML disease feature. Consistent with the prevailing view that MDS originates from the HSC compartment^57^, SEPTIN7 mis-splicing was confined to CD45^+^CD34^+^ HSPCs and absent from CD45^+^CD34^-^mature hematopoietic cells. The presence of the SEPTIN7 A3SS isoform was independent of *DDX41*-R525H VAF, indicating that aberrant splicing tracks with the presence of the somatic allele independent of the R525H clone size in whole bone marrow of patients with *DDX41* mutations.

Mechanistically, the A3SS event in SEPTIN7 introduces a premature stop codon, resulting in a selective and post-transcriptional reduction in SEPTIN7 protein in DDX41-depleted AML cells expressing the R525H or G530D somatic variant, but not in cells expressing germline variants or alanine substituted mutants. Septins are highly conserved GTP-binding cytoskeletal proteins involved in cytoskeleton organization, cytokinesis, cell polarity and membrane dynamics^52^. SEPTIN7 is the non-redundant, central subunit of the Septin core complex that stabilizes hetero-hexameric filaments with SEPTIN6 and SETPIN2^52,58^. Studies have shown that SEPTIN6 and SEPTIN2 protein levels are decreased on SEPTIN7 deletion in mouse fibroblasts^53^. Consistent with this finding, re-expression of R525H or G530D in our isogenic AML cells co-depleted SEPTIN6 at the protein level, while SEPTIN2 levels remained unchanged. We predict that partial loss of SEPTIN7 in *DDX41* biallelic cells would therefore destabilize the septin ring and deregulate cytokinesis and cell polarity. Relevant to hematopoiesis, *Septin7*-deficient HSCs lose cell polarity and show impaired proliferation and differentiation potential^35^ and *Septin7* was critical for early hematopoiesis in mice^36^. Further, a germline mutation in *SEPTIN6* causes pediatric MDS with severe neutropenia, morphologic dysplasia and aneuploidy^59^, and re-analysis of published RNAseq datasets^60^ revealed that SEPTIN6 is among the most commonly mis-spliced genes in *SF3B1*-mutant MDS. These studies indicate an important role for SEPTINs in hematopoiesis and myeloid neoplasms. SEPTIN7 is one of many mis-spliced targets of DDX41 R525H, and its causal role in impaired hematopoiesis and bone marrow failure remains to be proven. Its mutation-specific mis-splicing profile in patient HSPCs, the resulting loss of protein, and its function in cell division make it a compelling candidate linking DDX41 mis-splicing to dysplastic hematopoiesis.

In summary, this is the first study to demonstrate a role for somatic *DDX41* mutations in splicing regulation and to define the downstream splicing abnormalities of the *DDX41* biallelic state. These findings position *DDX41* alongside classical splicing factors – *SF3B1*, *SRSF2*, *U2AF1* and *ZRSR2* – that arise early and clonally in MDS, leading to inefficient hematopoietic differentiation. Aberrant splicing has emerged as a critical driver of HSC dysfunction across a spectrum of clonal myeloid diseases, and our findings in *DDX41* biallelic state is another clear example of this conceptual framework. Although the functional contribution of individual mis-splicing events to *DDX41*-mutant MDS remains to be formally determined, establishing the causative events is the necessary first step toward developing novel targeted therapies capable of eliminating or correcting the underlying defects for the treatment of bone marrow failure diseases.

## ACKNOWLEDGEMENTS

We thank members of the Lee lab for helpful discussion and critique of this manuscript; Tim Monahan, Bella Morocho, Grace Whitten and the Fred Hutch Cancer Center/University of Washington Hematopoietic Diseases Repository (FHCC/UW-HDR) for MDS and AML biospecimens; Hans-Peter Kiem for the pRSC32 lentiviral vector. R. Venkataraman is supported by the American Society of Hematology Graduate Hematology Award. S. Sinha is supported by the American Cancer Society Postdoctoral Fellowship (PF-23-1145387-01-ET). R.S. Welner is supported by the National Heart, Lung, and Blood Institute (P01 HL131477), a Mark Foundation for Cancer Research Endeavor Award, and an Edward P. Evans Foundation Discovery Research Grant. R. Lu is supported by grants from National Cancer Institute (R01 CA259480), American Cancer Society (RSG-22-036-01-DMC), Gabrielle’s Angel Foundation for Cancer Research and a Mark Foundation for Cancer Research Endeavor Award. P. B. Ferrell is supported by a Mark Foundation for Cancer Research Endeavor Award, a 350 Novartis Global Scholar Award, the National Institute of Diabetes and Digestive and Kidney Diseases (R56 DK138826), and VA MERIT Award 351 (I01BX005991). S.M. Fica is supported by a Wellcome Trust and Royal Society Sir Henry Dale Fellowship (ALR02270). S.C. Lee is supported by grants from National Cancer Institute (R01 CA292932), National Institute of Diabetes and Digestive and Kidney Diseases (RC2 DK127989), an Edward P. Evans Foundation Discovery Research Grant, and a Mark Foundation for Cancer Research Endeavor Award. This research is also supported by the Fred Hutchinson Cancer Center Shared Resources through NCI Cancer Center Support Grants (P30 CA015704 and S10 OD028685).

## Author Contributions

R.V. designed research, performed research, analyzed data, and prepared the manuscript. N.P., T-Y.H, E.A.B., F-Y.C., E.A.A-G., A.S., T.M., J.R., O.A., S.S., E.E.M., X.C., K.K., and S.M.F. performed research and analyzed data. M.P.C. J.S.A., D.L.S. and A.R. provided vital reagents. R.S.W., R.L., O.A-W., and P.B.F., provided critical feedback for the study. S.C.L. designed research, performed research, analyzed data, prepared the manuscript, supervised research and secured research funding. All authors have read, reviewed and consented to submission of this manuscript.

## REFERENCES

1 Lewinsohn, M. et al. Novel germ line DDX41 mutations define families with a lower age of MDS/AML onset and lymphoid malignancies. Blood 127, 1017–1023 (2016). 10.1182/blood-2015-10-676098

2 Makishima, H. et al. Germ line DDX41 mutations define a unique subtype of myeloid neoplasms. Blood 141, 534–549 (2023). 10.1182/blood.2022018221

3 Polprasert, C. et al. Inherited and Somatic Defects in DDX41 in Myeloid Neoplasms. Cancer Cell 27, 658–670 (2015). 10.1016/j.ccell.2015.03.017

4 Feurstein, S. et al. Germline variants drive myelodysplastic syndrome in young adults. Leukemia 35, 2439–2444 (2021). 10.1038/s41375-021-01137-0

5 Duployez, N. et al. Prognostic impact of DDX41 germline mutations in intensively treated acute myeloid leukemia patients: an ALFA-FILO study. Blood 140, 756–768 (2022). 10.1182/blood.2021015328

6 Li, P. et al. AML with germline DDX41 variants is a clinicopathologically distinct entity with an indolent clinical course and favorable outcome. Leukemia 36, 664–674 (2022). 10.1038/s41375-021-01404-0

7 Kadono, M. et al. Biological implications of somatic DDX41 p.R525H mutation in acute myeloid leukemia. Exp Hematol 44, 745–754 e744 (2016). 10.1016/j.exphem.2016.04.017

8 Badar, T. & Chlon, T. Germline and Somatic Defects in DDX41 and its Impact on Myeloid Neoplasms. Curr Hematol Malig Rep 17, 113–120 (2022). 10.1007/s11899-022-00667-3

9 Makishima, H., Bowman, T. V. & Godley, L. A. DDX41-associated susceptibility to myeloid neoplasms. Blood 141, 1544–1552 (2023). 10.1182/blood.2022017715

10 Zhang, Z. et al. The helicase DDX41 senses intracellular DNA mediated by the adaptor STING in dendritic cells. Nat Immunol 12, 959–965 (2011). 10.1038/ni.2091

11 Parvatiyar, K. et al. The helicase DDX41 recognizes the bacterial secondary messengers cyclic di-GMP and cyclic di-AMP to activate a type I interferon immune response. Nat Immunol 13, 1155–1161 (2012). 10.1038/ni.2460

12 Lee, K. G. et al. Bruton’s tyrosine kinase phosphorylates DDX41 and activates its binding of dsDNA and STING to initiate type 1 interferon response. Cell Rep 10, 1055–1065 (2015). 10.1016/j.celrep.2015.01.039

13 Weinreb, J. T. et al. Excessive R-loops trigger an inflammatory cascade leading to increased HSPC production. Dev Cell 56, 627–640 e625 (2021). 10.1016/j.devcel.2021.02.006

14 Chlon, T. M. et al. Germline DDX41 mutations cause ineffective hematopoiesis and myelodysplasia. Cell Stem Cell 28, 1966–1981 e1966 (2021). 10.1016/j.stem.2021.08.004

15 Tungalag, S. et al. Ribosome profiling analysis reveals the roles of DDX41 in translational regulation. Int J Hematol 117, 876–888 (2023). 10.1007/s12185-023-03558-2

16 Mosler, T. et al. R-loop proximity proteomics identifies a role of DDX41 in transcription-associated genomic instability. Nat Commun 12, 7314 (2021). 10.1038/s41467-021-27530-y

17 Shinriki, S. et al. DDX41 coordinates RNA splicing and transcriptional elongation to prevent DNA replication stress in hematopoietic cells. Leukemia 36, 2605–2620 (2022). 10.1038/s41375-022-01708-9

18 Bi, H. et al. DDX41 resolves G-quadruplexes to maintain erythroid genome integrity and prevent cGAS-mediated cell death. Nat Commun 16, 7195 (2025). 10.1038/s41467-025-62307-7

19 Dybkov, O. et al. Regulation of 3’ splice site selection after step 1 of splicing by spliceosomal C* proteins. Sci Adv 9, eadf1785 (2023). 10.1126/sciadv.adf1785

20 Osterhoudt, K., Bagno, O., Katzman, S. & Zahler, A. M. Spliceosomal helicases DDX41/SACY-1 and PRP22/MOG-5 both contribute to proofreading against proximal 3’ splice site usage. RNA 30, 404–417 (2024). 10.1261/rna.079888.123

21 Kim, J. A. et al. Oncogenic DEAD-box ATPase DDX41 establishes transcript ensembles via CLK3-dependent and -independent mechanisms. Nat Commun 16, 9716 (2025). 10.1038/s41467-025-65195-z

22 Ma, J., Mahmud, N., Bosland, M. C. & Ross, S. R. DDX41 is needed for pre-and postnatal hematopoietic stem cell differentiation in mice. Stem Cell Reports 17, 879–893 (2022). 10.1016/j.stemcr.2022.02.010

23 Stepanchick, E. et al. DDX41 haploinsufficiency causes inefficient hematopoiesis under stress and cooperates with p53 mutations to cause hematologic malignancy. Leukemia 38, 1787–1798 (2024). 10.1038/s41375-024-02304-9

24 Omura, H. et al. Structural and Functional Analysis of DDX41: a bispecific immune receptor for DNA and cyclic dinucleotide. Sci Rep 6, 34756 (2016). 10.1038/srep34756

25 Cordin, O., Hahn, D. & Beggs, J. D. Structure, function and regulation of spliceosomal RNA helicases. Curr Opin Cell Biol 24, 431–438 (2012). 10.1016/j.ceb.2012.03.004

26 Jurica, M. S., Licklider, L. J., Gygi, S. R., Grigorieff, N. & Moore, M. J. Purification and characterization of native spliceosomes suitable for three-dimensional structural analysis. RNA 8, 426–439 (2002). 10.1017/s1355838202021088

27 Yoshida, K. et al. Frequent pathway mutations of splicing machinery in myelodysplasia. Nature 478, 64–69 (2011). 10.1038/nature10496

28 Thol, F. et al. Frequency and prognostic impact of mutations in SRSF2, U2AF1, and ZRSR2 in patients with myelodysplastic syndromes. Blood 119, 3578–3584 (2012). 10.1182/blood-2011-12-399337

29 Pelossof, R. et al. Prediction of potent shRNAs with a sequential classification algorithm. Nat Biotechnol 35, 350–353 (2017). 10.1038/nbt.3807

30 Fellmann, C. et al. An optimized microRNA backbone for effective single-copy RNAi. Cell Rep 5, 1704–1713 (2013). 10.1016/j.celrep.2013.11.020

31 Van Nostrand, E. L. et al. Robust, Cost-Effective Profiling of RNA Binding Protein Targets with Single-end Enhanced Crosslinking and Immunoprecipitation (seCLIP). Methods Mol Biol 1648, 177–200 (2017). 10.1007/978-1-4939-7204-3_14

32 Mars, Z. et al. Biallelic germline variants in the hematologic malignancy predisposition gene DDX41 cause retinal dystrophy through dysregulation of retinal homeostasis. medRxiv (2026). 10.64898/2026.01.28.26344834

33 Fica, S. M., Oubridge, C., Wilkinson, M. E., Newman, A. J. & Nagai, K. A human postcatalytic spliceosome structure reveals essential roles of metazoan factors for exon ligation. Science 363, 710–714 (2019). 10.1126/science.aaw5569

34 Cesana, M. et al. A CLK3-HMGA2 Alternative Splicing Axis Impacts Human Hematopoietic Stem Cell Molecular Identity throughout Development. Cell Stem Cell 22, 575–588 e577 (2018). 10.1016/j.stem.2018.03.012

35 Kandi, R. et al. Cdc42-Borg4-Septin7 axis regulates HSC polarity and function. EMBO Rep 22, e52931 (2021). 10.15252/embr.202152931

36 Ronkina, N. et al. Septin7 is essential in early hematopoiesis, but redundant at later stages. Life Sci Alliance 9 (2026). 10.26508/lsa.202603637

37 Hong, Y. G. et al. The RNA m6A Reader YTHDF1 Is Required for Acute Myeloid Leukemia Progression. Cancer Res 83, 845–860 (2023). 10.1158/0008-5472.CAN-21-4249

38 Hwang, W. C. et al. Impaired binding affinity of YTHDC1 with METTL3/METTL14 results in R-loop accumulation in myelodysplastic neoplasms with DDX41 mutation. Leukemia 38, 1353–1364 (2024). 10.1038/s41375-024-02228-4

39 Luis, T. C., Ichii, M., Brugman, M. H., Kincade, P. & Staal, F. J. Wnt signaling strength regulates normal hematopoiesis and its deregulation is involved in leukemia development. Leukemia 26, 414–421 (2012). 10.1038/leu.2011.387

40 Caliskan, C., Yuce, Z. & Ogun Sercan, H. Dvl proteins regulate SMAD1, AHR, mTOR, BRD7 protein expression while differentially regulating canonical and non-canonical Wnt signaling pathways in CML cell lines. Gene 854, 147109 (2023). 10.1016/j.gene.2022.147109

41 Linder, P. & Jankowsky, E. From unwinding to clamping - the DEAD box RNA helicase family. Nat Rev Mol Cell Biol 12, 505–516 (2011). 10.1038/nrm3154

42 Mallam, A. L., Del Campo, M., Gilman, B., Sidote, D. J. & Lambowitz, A. M. Structural basis for RNA-duplex recognition and unwinding by the DEAD-box helicase Mss116p. Nature 490, 121–125 (2012). 10.1038/nature11402

43 Gilman, B., Tijerina, P. & Russell, R. Distinct RNA-unwinding mechanisms of DEAD-box and DEAH-box RNA helicase proteins in remodeling structured RNAs and RNPs. Biochem Soc Trans 45, 1313–1321 (2017). 10.1042/BST20170095

44 Stavrou, S., Aguilera, A. N., Blouch, K. & Ross, S. R. DDX41 Recognizes RNA/DNA Retroviral Reverse Transcripts and Is Critical for In Vivo Control of Murine Leukemia Virus Infection. mBio 9 (2018). 10.1128/mBio.00923-18

45 Kuhn, R., Schwenk, F., Aguet, M. & Rajewsky, K. Inducible gene targeting in mice. Science 269, 1427–1429 (1995). 10.1126/science.7660125

46 Lee, S. C. et al. Synthetic Lethal and Convergent Biological Effects of Cancer-Associated Spliceosomal Gene Mutations. Cancer Cell 34, 225–241 e228 (2018). 10.1016/j.ccell.2018.07.003

47 Smeets, M. F. et al. The Rothmund-Thomson syndrome helicase RECQL4 is essential for hematopoiesis. J Clin Invest 124, 3551–3565 (2014). 10.1172/JCI75334

48 Zhu, X. et al. Identification of functional cooperative mutations of SETD2 in human acute leukemia. Nat Genet 46, 287–293 (2014). 10.1038/ng.2894

49 Zhang, Y. L. et al. Setd2 deficiency impairs hematopoietic stem cell self-renewal and causes malignant transformation. Cell Res 28, 476–490 (2018). 10.1038/s41422-018-0015-9

50 Shibata, S. et al. SLF2 and SMC5 dysfunction drives HSC aging and predisposes to MDS, defining a new inherited bone marrow failure syndrome. Leukemia (2026). 10.1038/s41375-026-03061-7

51 Gallardo, M. et al. hnRNP K Is a Haploinsufficient Tumor Suppressor that Regulates Proliferation and Differentiation Programs in Hematologic Malignancies. Cancer Cell 28, 486–499 (2015). 10.1016/j.ccell.2015.09.001

52 Neubauer, K. & Zieger, B. The Mammalian Septin Interactome. Front Cell Dev Biol 5, 3 (2017). 10.3389/fcell.2017.00003

53 Menon, M. B. et al. Genetic deletion of SEPT7 reveals a cell type-specific role of septins in microtubule destabilization for the completion of cytokinesis. PLoS Genet 10, e1004558 (2014). 10.1371/journal.pgen.1004558

54 Fisher, J. S. et al. Functional Characterization of Myeloid Neoplasm-associated DDX41 Variants Reveals Pathogenic Interaction with Acquired Hotspot Mutation. bioRxiv, 2026.2005.2027.727893 (2026). 10.64898/2026.05.27.727893

55 Liu, F., Putnam, A. A. & Jankowsky, E. DEAD-box helicases form nucleotide-dependent, long-lived complexes with RNA. Biochemistry 53, 423–433 (2014). 10.1021/bi401540q

56 Bohnsack, K. E., Yi, S., Venus, S., Jankowsky, E. & Bohnsack, M. T. Cellular functions of eukaryotic RNA helicases and their links to human diseases. Nat Rev Mol Cell Biol 24, 749–769 (2023). 10.1038/s41580-023-00628-5

57 Nimer, S. D. Myelodysplastic syndromes. Blood 111, 4841–4851 (2008). 10.1182/blood-2007-08-078139

58 Mendonca, D. C. et al. A revised order of subunits in mammalian septin complexes. Cytoskeleton (Hoboken) 76, 457–466 (2019). 10.1002/cm.21569

59 Renella, R. et al. Congenital X-linked neutropenia with myelodysplasia and somatic tetraploidy due to a germline mutation in SEPT6. Am J Hematol 97, 18–29 (2022). 10.1002/ajh.26382

60 Obeng, E. A. et al. Physiologic Expression of Sf3b1(K700E) Causes Impaired Erythropoiesis, Aberrant Splicing, and Sensitivity to Therapeutic Spliceosome Modulation. Cancer Cell 30, 404–417 (2016). 10.1016/j.ccell.2016.08.006

