## Supplemental Materials for "Somatic *DDX41* mutations confer neomorphic splicing activity in the pathogenesis of myelodysplastic neoplasms"

**SUPPLEMENTAL FIGURE LEGENDS**

**Figure S1, related to Fig 1: (A)** Western blot of MOLM-13 cells expressing doxycycline inducible shRNA targeting DDX41 3'UTR at 72h with and without doxycycline. Cell proliferation curves in MOLM-13 isogenic cells overexpressing **(B)** germline DD4X1 mutants, **(C)** somatic DDX41 mutants, and **(D)** structural DDX41 mutants in the absence of doxycycline.

**Figure S2, related to Fig 2: (A)** Inclusion level scatter plots showing percent spliced-in of DDX41-wildtype (Y-axis) against percent spliced-in (PSI) of DDX41 mutants (X-axis). Blue dots denote events with FDR < 0.05 and delta-PSI > 0.2 (promoted in DDX41 mutants), and red dots denote events with FDR < 0.05 and delta-PSI < 0.2 (repressed in DDX41 mutants). **(B)** Bubble plots summarizing significant splicing events across DDX41 germline mutants (G173R, E256K, Y259C and S363del) by splicing event type (A3SS, A5SS, RI, SE). Bubble size represents the number of significant splicing events and color represents the median delta-PSI for that event category. Significant splicing events ranked by delta-PSI > 0.2 and FDR <0.01, promoted in DDX41 germline mutant **(C)** G173R, **(D)** E256K, **(E)** Y259C and **(F)** S363del. **(G)** IGV coverage plots of additional statistically significant mis-spliced events: retained intron in MED25, A3SS usage in YTHDF1, retained intrin in FGD2 and A3SS usage in SMC5, Y-axis, RPMK normalized coverage. **(H)** Principal Component Analysis (PCA) plot for RNA-seq samples based on gene expression across DDX41 somatic and germline mutants, DDX41 wildtype and empty vector controls, colored by condition. **(I)** Unsupervised hierarchical clustering heatmap for the union of differentially expressed genes identified across DDX41-mutant versus wild-type comparisons. Rows represent genes and columns represent DDX41 mutants in triplicate. **(J)** Unsupervised hierarchical clustering heatmap for the intersection of differentially expressed genes identified for DDX41-R525H and DDX41-G530D versus wild-type comparisons. Rows represent genes and columns represent DDX41 mutant in triplicate.

**Figure S3, related to Fig 3: (A)** Metagene plot of average eCLIP peak density across length-scaled transcripts (5′ UTR, CDS, 3′ UTR) for DDX41-WT (yellow) and R525H (magenta). **(B)** GO analysis among R525H-bound genes, ranked by $-\log_{10}$(p-value), performed with Metascape. **(C)** AlphaFold model of DDX41 with the RNA path and ATP modeled from Dbp5 (left) and the same model rotated 45° (right), showing the positions of the leukemia-associated residues R525H, G530D, and E256K relative to the Q motif, DEAD box, HELIC domain, and $Zn^{++}$ knuckle.

**Figure S4, related to Fig 4: (A)** Live cell number over 7 days in culture for cells not treated with doxycycline, and expressing empty vector, DDX41-WT, R525A, or G530A (data are mean ± SD of n=3).

**Figure S5, related to Fig 5: (A)** Schematic of validation of the germline *Ddx41*$^{KO/+}$ mice. **(B)** Schematic of Cre-mediated recombination converting the floxed *Ddx41* allele to the R525H knock-in allele (GCA→GTA, exon 15). Sanger sequencing of bone marrow confirms the heterozygous mutation in *Ddx41*$^{R525H/+}$ mice versus wild-type control. **(C)** GO analysis among genes mis-spliced in biallelic *Ddx41*$^{KOI/R525H}$ mice, ranked by $-\log_{10}$ (p-value), analysis performed with Metascape. **(D)** Coverage tracks showing alternative 3′ splice site (A3SS) usage in *Setd2 and* retained intron in *Clk3* for the mentioned genotypes.

**Figure S6, related to Fig 6: (A)** Mutational status and variant allele frequency (VAF) of *DDX41* mutant patient samples. **(B)** Alignment of the mouse and human SEPTIN7 3’ intron 1/exon 2 region which is the 144 base inclusion on mis-splicing on SEPTIN7; asterisks mark conserved bases, the boxed sequence is exon 2, and the highlighted bases indicate the canonical 3′ splice site. **(C)** SEPTIN7 western blot in DDX41-knockdown cells reconstituted with empty vector (EV), DDX41-WT, R525H, G530D, R525A, or G530A; GAPDH, loading control. **(D)** Quantification of SEPTIN7 relative to GAPDH from (C).

### Supplemental Figure S1

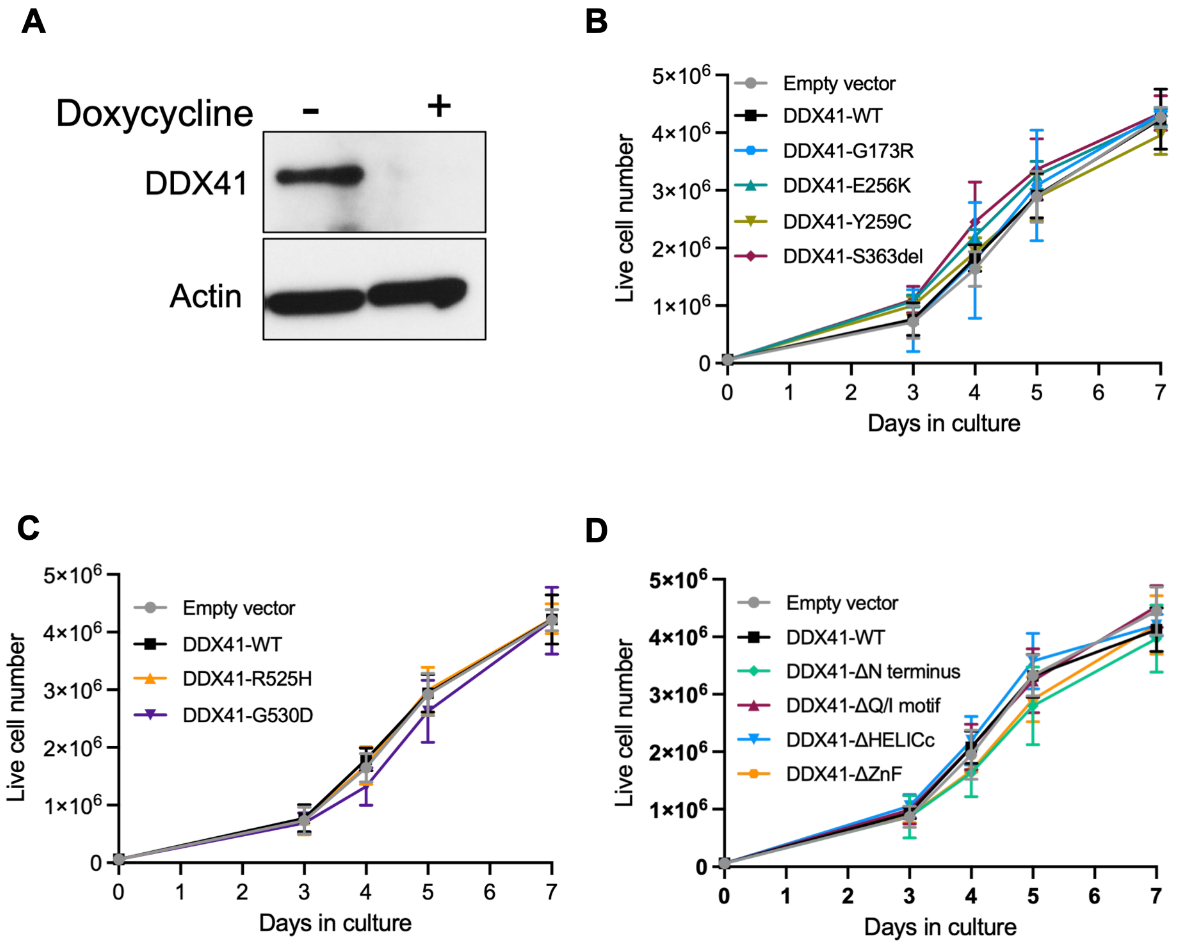

### Supplemental Figure S2

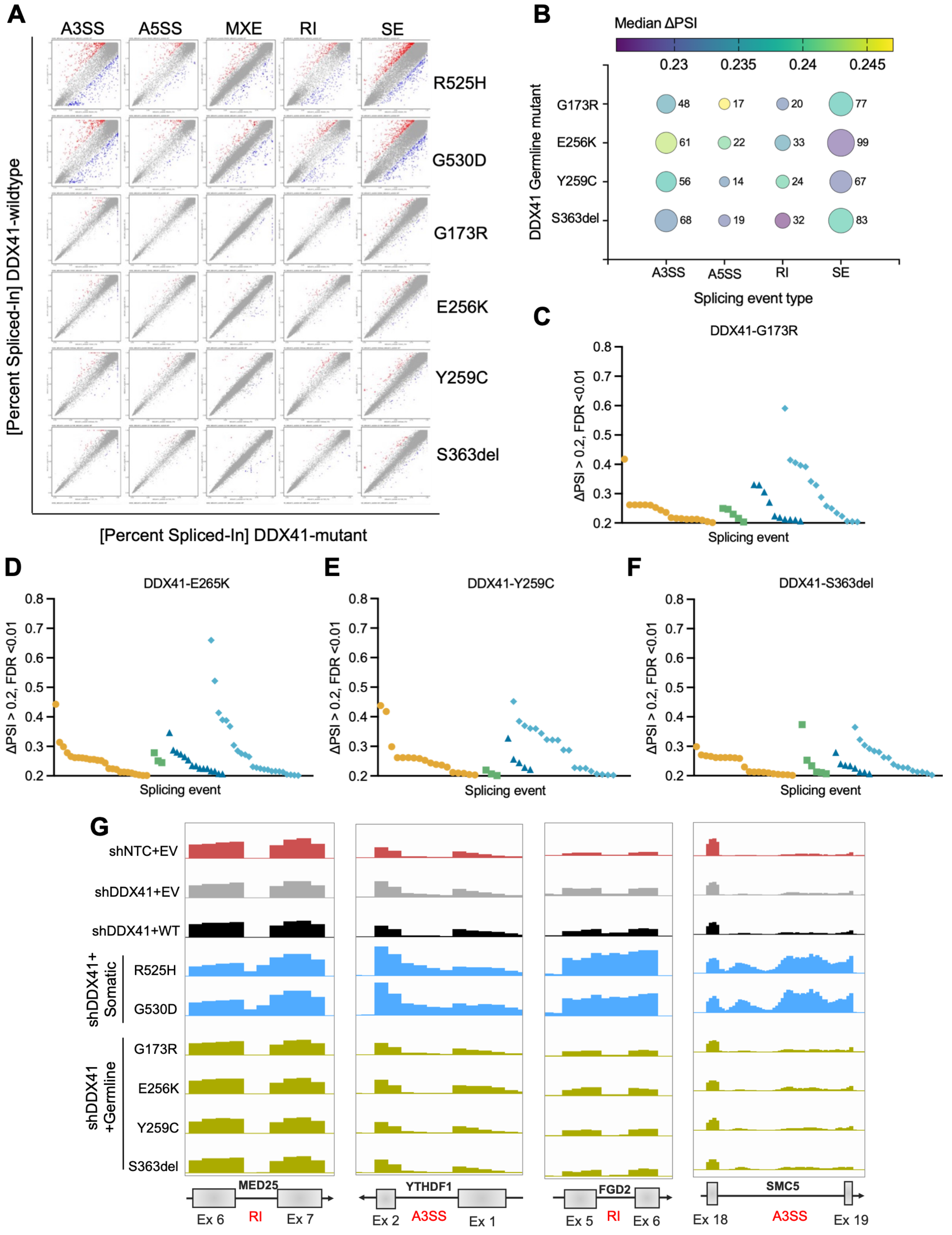

### Supplemental Figure S2, Contd.

H

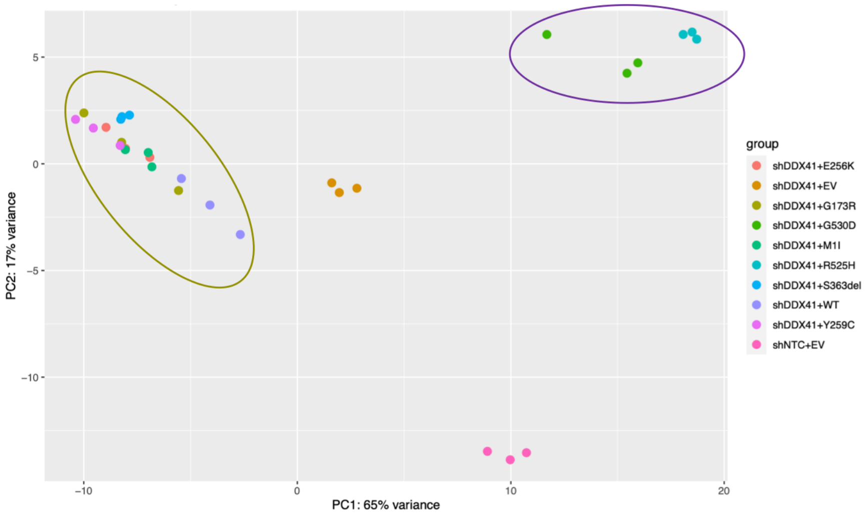


I

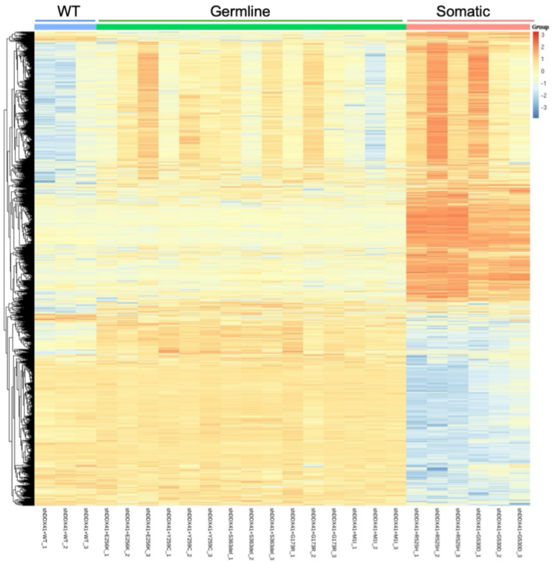


J

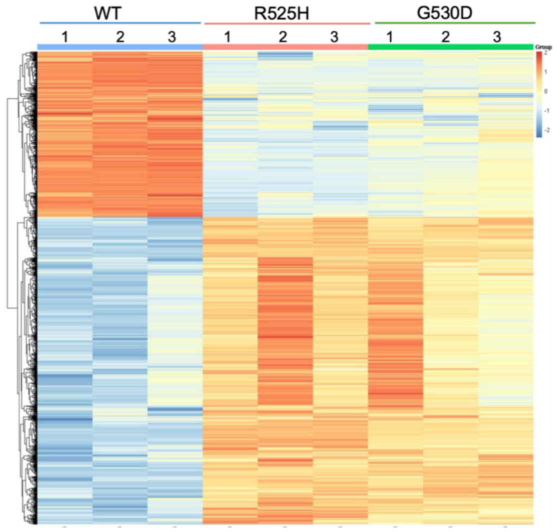

### Supplemental Figure S3

A

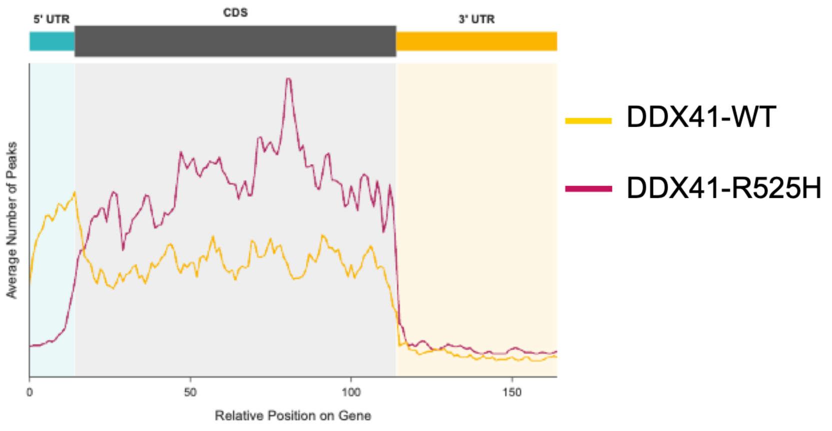


B

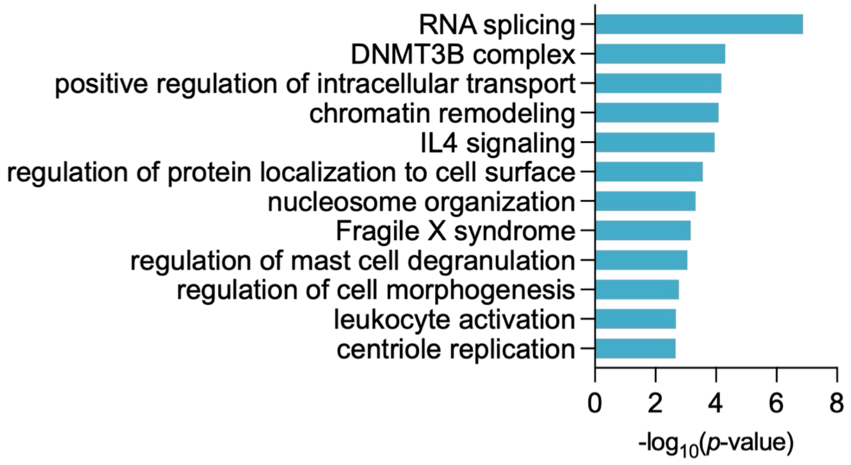


C

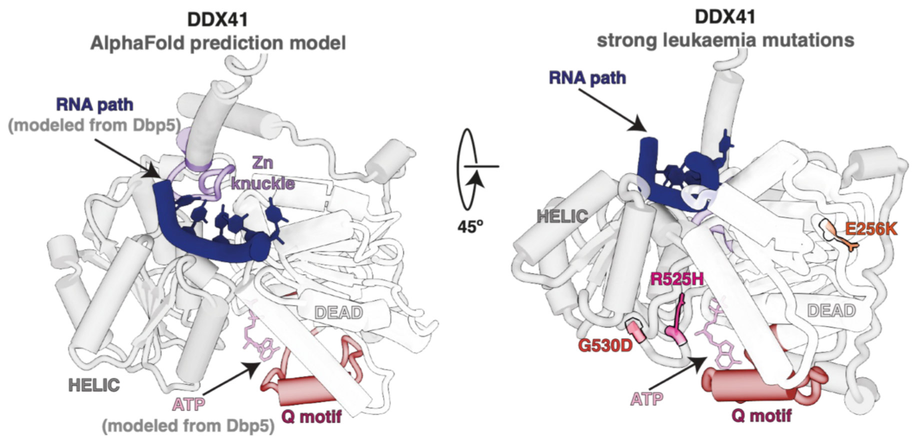

### Supplemental Figure S4

**A**

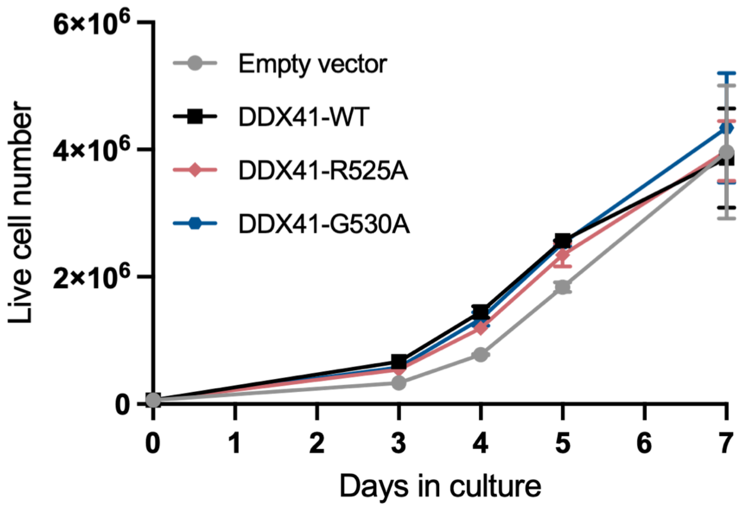

### Supplemental Figure S5

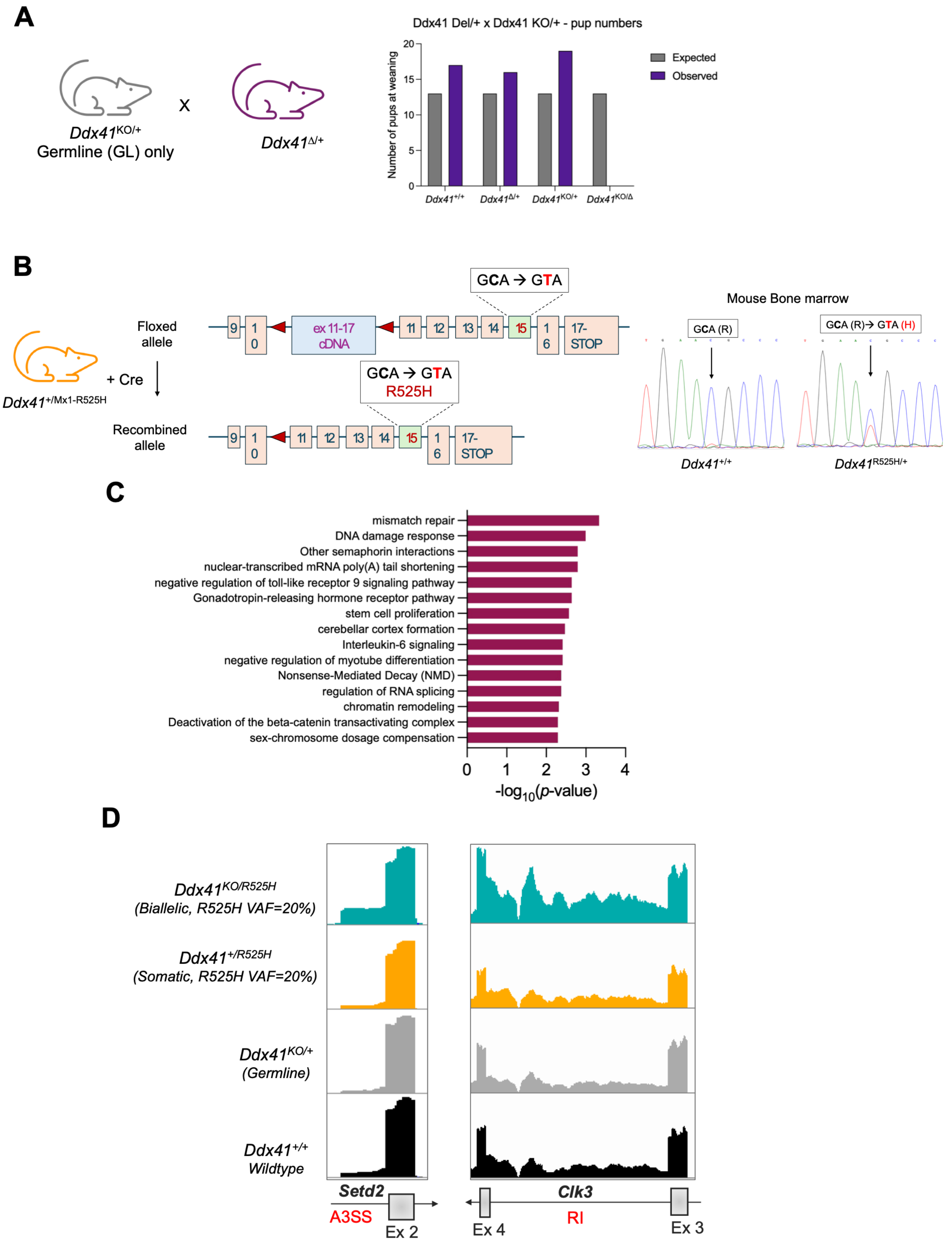

### Supplemental Figure S6

### A

| Patinet ID | Diagnosis | Prior BM Transplant | *DDX41* germline mutation | *DDX41* germline mutation VAF | *DDX41* somatic mutation | *DDX41* somatic mutation VAF | Other mutations |
|---|---|---|---|---|---|---|---|
| Pt_001 | AML | No | Wildtype | n/a | Wildtype | n/a | *ASXL1, EZH2, FLT3, NRAS, TET2* |
| Pt_002 | AML | No | Wildtype | n/a | Wildtype | n/a | *ASXL1, CEBPA, EZH2, FLT3, SRSF2, STAG2, TET2* |
| Pt_003 | AML | No | Wildtype | n/a | Wildtype | n/a | *ASXL1, EZH2, FLT3, PTPN11, TP53* |
| Pt_004 | AML | No | c.27+16G>C | 50% | none | n/a | *ASXL1, BCPR, CSF3R, EZH2, FLT3, RUNX1* |
| Pt_005 | MDS | No | p.A191T | 49% | none | n/a | none |
| Pt_006 | AML | No | c.1098+6G>A | 50% | none | n/a | *ASXL1, EZH2, FLT3, KRAS, MPL, NRAS* |
| Pt_007 | AML | No | p.M1I? | 50% | p.R525H | 3% | none |
| Pt_008 | MDS | No | p.E256K | 45% | p.R525H | 28% | none |
| Pt_009 | AML | Yes | p.D140Gfs*2 | 10% | p.R525H | 11% | *TP53* |
| Pt_010 | AML | No | p.K102Rfs*32 | 48% | p.R525H | 9% | *PHF6, EZH2* |
| Pt_011 | AML | No | p.F242_R249del | 11% | p.R525H | 14% | *CARD11, GNAS, PHF6* |
| Pt_012 | AML | No | p.M1I? | 48% | p.R525H | 4% | *SF3B1, FANCC* |

### B

```
Mouse GCTATGCTGCAATGGATCTGCTACTCCTGTTTGGACTGAATGCGCCATTACAATT--TGG
Human ---GTGTTTCAATGGATTTGCCTCTCCTGTTTGGACTGAATATGTCATTACAATTCTTGG
          ** * ******** ***  ******************  * **********  ***

Mouse AATTCTTCACTTGCTGT-CTTCTGTGAATTTGTTCTGTTGGAATCTGGAACATTGGAATG
Human AATTCTTTGTTTACTGTTCCTCTGTGGATTTCTTCTGTTGGAATCTGGAACACTGGAATG
      *******   ** **** * ****** **** ******************** *******

Mouse ATGGAAATGTTTCCATTACAAAGTAGCT
Human ATGGAAATGTGTTCTTTACAAAGTAGCT
      ********** * * *************
                         Ex 2
```

### C

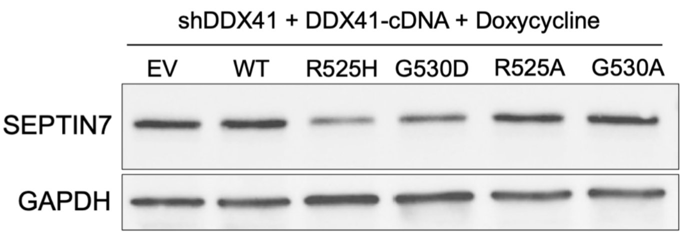


### D

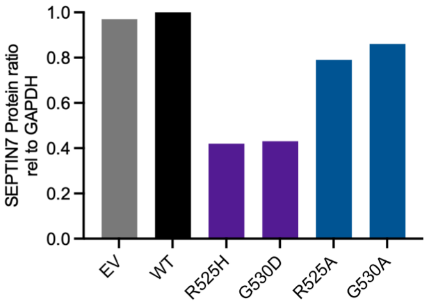
